# Neuroanatomical characterization of the Nmu-Cre knock-in mice reveals an interconnected network of unique neuropeptidergic cells

**DOI:** 10.1101/2023.01.19.524191

**Authors:** Mireia Medrano, Wissal Allaoui, Mathias Van Bulck, Sofie Thys, Leila Makrini-Maleville, Eve Seuntjens, Winnok H. De Vos, Emmanuel Valjent, Bálazs Gaszner, Ann Van Eeckhaut, Ilse Smolders, Dimitri De Bundel

## Abstract

Neuromedin U (NMU) is an evolutionary conserved neuropeptide that has been implicated in multiple processes, such as circadian regulation, energy homeostasis, reward processing and stress coping. Although central expression of NMU has been addressed previously, the lack of specific and sensitive tools has prevented a comprehensive characterization of NMU-expressing neurons in the brain. We have generated a knock-in mouse model constitutively expressing Cre recombinase under the *Nmu* promoter. We have validated the model using a multi-level approach based on quantitative reverse-transcription polymerase chain reactions, *in situ* hybridization, a reporter mouse line and an adenoviral vector driving Cre-dependent expression of a fluorescent protein. Using the Nmu-Cre mouse, we performed a complete mapping of NMU expression in adult mouse brain, unveiling a potential midline NMU modulatory circuit with the ventromedial hypothalamic nucleus (VMH) as a key node. Moreover, immunohistochemical analysis suggested that NMU neurons in the VMH mainly constitute a unique population of hypothalamic cells. Taken together, our results suggest that Cre expression in the Nmu-Cre mouse model largely reflects NMU expression in the adult mouse brain, without altering endogenous NMU expression. Thus, the Nmu-Cre mouse model is a powerful and sensitive tool to explore the role of NMU neurons in mice.

## 1. Introduction

Neuromedin U (NMU) is recognized as a multifunctional neuropeptide involved in various physiological processes, such as circadian regulation, energy homeostasis and regulation of feeding behavior, reproductive behaviors, reward processing and stress coping^1–4^. However, knowledge on the distribution, expression and connectivity of NMU-producing cells in the brain remains limited. NMU is coded by the *Nmu* gene and shows a notable amino acid sequence homology across different species^3^. The high degree of amino acid sequence conservation is indicative of a strong evolutionary pressure to maintain its structure and function across not only mammals but the entire vertebrate subphylum^1–5^. Moreover, NMU analogs are conserved across bilaterian animals, including invertebrates^6–8^. In mammals, NMU exists mainly in two principal forms, a long amino acid peptide (NMU-23 in rodents, NMU-25 in humans) and a truncated peptide (NMU-8), although other forms have also been reported^2,9^. The asparagine-linked amidation at the C-terminus of NMU is an important structural characteristic of the peptide. Indeed, this posttranslational modification has shown to be indispensable for receptor binding and activation^10–13^. NMU exerts its biological effects via two known G protein-coupled receptors: NMUR1 which is mainly expressed in the periphery and NMUR2, principally expressed in the central nervous system (CNS)^1^.

NMU distribution in the CNS has been investigated in different species including human, rat, and to a lesser extent in mouse. These studies used chromatographic techniques, radioimmunoassays (RIA) and immunohistochemical (IHC) analyses to detect NMU-like immunoreactivity (NMU-LI)^14–22^, and *in situ* hybridization (ISH), quantitative reverse-transcription polymerase chain reaction (RT-qPCR) and Northern blot analysis to study *Nmu* mRNA distribution^10,11,15,23–32^. Compared to expression levels in the small intestine, low or moderate levels of NMU-LI have been reported in the CNS, both in the brain and the spinal cord^1,3^. Within the brain, the highest levels of NMU-LI were detected in rat, human and mouse pituitary gland^16,19,22^. Moderate to high levels of *Nmu* mRNA and NMU-LI where found in the hypothalamus^16,19,21,25,26,32^, with highest levels in the mouse suprachiasmatic nucleus (SCh), dorsomedial hypothalamic nucleus (DMH), arcuate nucleus (ARC) and ventromedial hypothalamic nucleus (VMH)^26^, supporting the described role of NMU in circadian regulation, reproduction, and feeding and energy homeostasis^1–4^. However, it should be noted that previous descriptions of NMU expression in the CNS shows a certain level of inconsistency. Histochemical techniques often have insufficient sensitivity to detect low-abundant molecules and the currently commercially available anti-NMU antibodies show low specificity. Furthermore, most IHC studies used primary antisera raised against synthetic porcine NMU-8, using colchicine to block axonal transport and increase the peptide content in cell bodies^16,20,22^. Potential cross-reactivity and inherent differences between antiserum batches could partially explain the lack of reproducible results regarding NMU distribution. Moreover, a previous RIA analysis showed interspecies molecular differences of NMU-LI^18^, pointing out that the NMU distribution reported in different species using only synthetic porcine NMU-8 should be analyzed with caution. In addition, colchicine-pretreatment can alter normal brain physiology, limiting its use for studying peptide distribution under normal conditions^33^. Moreover, classical ISH techniques lack the sensitivity required to detect many low-abundant transcripts^34,35^. Indeed, a few studies have addressed *Nmu* mRNA expression by ISH in mice, showing relevant discrepancies in terms of level of expression and distribution^10,21,26,36^. Therefore, a more detailed and unambiguous description of the distribution, expression and connectivity of NMU-producing cells in the brain would enhance our understanding on the central role of NMU in health and disease.

Over the past two decades, technical developments have created opportunities to access neural systems at a molecular, cellular, circuit and functional level^37,38^. Bacterial artificial chromosome (BAC)-mediated transgenics and gene targeting are approaches that are widely used to drive the expression of specific genes of interest^38,39^. One of the most extensively used cell-type-specific genetic targeting strategies is the binary system Cre/LoxP. In this system, the bacteriophage recombinase Cre transgene, of which the expression is under the regional control of specific gene promotors, controls the expression of a target gene by inducing the recombination between two LoxP sites^38,40,41^. Mice that express Cre under a cell-type-specific promoter are instrumental for functional neuroanatomical characterization. Cre drivers can be cross-bred with reporter lines that carry a fluorescent protein to provide cell-specific expression of the fluorescent marker. Moreover, the Cre driver can be targeted with Cre-dependent viral vectors to manipulate cells of interest in a temporal and regional-specific manner. This permits the elucidation of the neuroanatomical organization and functions of the targeted cells. At present, two BAC transgenic mice have been generated driving the Cre recombinase gene or the enhanced green fluorescent protein (EGFP) reporter gene under the *Nmu* promoter but they have not yet been systematically validated^42^. In addition, it should be noted that BAC transgenes are generated by nonspecific integration of the transgene into the target genome. Therefore, the transgene can be inserted into an unknown locus in the genome, leading to ectopic expression and affecting expression of neighbouring genes^43^. Moreover, expression variegation makes it necessary to functionally screen founder lines^43,44^, since the positional effect may cause a variable number of copies to be incorporated into the genome, thus affecting the level of Cre expression^38,43–45^.

In the present study, we generated and validated a novel Nmu-Cre knock-in mouse model constitutively expressing Cre recombinase under the endogenous *Nmu* promoter. The validation was ensured by cross-breeding the Nmu-Cre mouse with a reporter line and by detecting *Nmu* mRNA expression using RT-qPCR and ISH. We performed a detailed neuroanatomical mapping of NMU-expressing cells in the mouse brain and characterized these cells using a viral tracer, IHC and ISH analysis.

## 2. Materials and methods

### 2.1. Animals

The B6.NmuCre-IRES-Nmu or Nmu-Cre knock-in mouse line was developed in collaboration with genOway (France) on a C57BL/6 genetic background as described below. PCR-selected male and female Nmu-Cre heterozygous mice were backcrossed to C57BL/6J mice (Janvier, France) to maintain the breeding colony. B6.Cg-*Gt(ROSA)26Sor^tm6(CAG-ZsGreen1)Hze^*/J, also known as Ai6 or Ai6(RCL-ZsGreen)^41^ mice (stock #007906, The Jackson Laboratory, USA; RRID IMSR_JAX:007906) were used as Cre reporter line. This line contains a loxP-flanked STOP cassette preventing transcription of a CAG promoter-driven enhanced green fluorescent protein variant (ZsGreen1). PCR-selected male and female Nmu-Cre heterozygous mice were crossed with homozygous Ai6 reporter mice to obtain Nmu-Cre:ZsGreen1 offspring that drive expression of the green fluorescent protein ZsGreen1 in a Cre-dependent manner. All mice were bred in-house at the animal facility of the Vrije Universiteit Brussel. All mice were adult (≥ 8 weeks) when set in breeding or at the start of experiments. Mice were group-housed (1290 eurostandard type III cages, Tecniplast, Italy) in a temperature (18–24°C) and humidity (30-70%) regulated environment with a 12/12h light/dark cycle (onset dark cycle: 6 p.m.). Mice had free access to food pellets (A03, SAFE, France) and water. Cages were minimally enriched with shelters, wooden gnawing blocks and nesting material. Nmu-Cre mice that underwent stereotaxic surgery were single housed afterwards (1264C Eurostandard type II cages, Tecniplast, Italy) for the remainder of the experiment. All mice used in the present study were sacrificed between 3 and 5 p.m. (light cycle). All experiments were executed by certified and experienced researchers and were approved by the Ethical Committee for Animal Experiments of the Faculty of Medicine and Pharmacy of the Vrije Universiteit Brussel. All experiments were performed according to the European Community Council directive (2010/63/EU) and the Belgium Royal Decree (29/05/2013), and complied with the ARRIVE guidelines^46^. All efforts were made to reduce stress and suffering of the animals to a minimum.

### 2.2. Generation of the Nmu-Cre knock-in mouse model

The selected targeting strategy for generation of the Nmu-Cre knock-in mouse model (Figure 1) consisted of the insertion of a Cre-IRES-*Nmu* cDNA in frame with exon 1 ATG of the *Nmu* gene on chromosome 5qC3.3, maintaining the expression of the most prevalent isoform encoding for the 174 amino acid precursor and minimizing both the potential risk of interfering with regulatory elements and the deregulation of the targeted and/or neighboring genes. The Cre-IRES-*Nmu* targeting vector was designed (from 5’ to 3’) with a Cre coding sequence, an internal ribosome entry site (IRES) element, the coding sequence of NMU isoform 1, an hGHpolyA signal and a neomycin cassette for positive selection, flanked by two flippase recognition target (FRT) sites (Figure 1B). To generate the targeting vector carrying the Cre cassette, two homology arms were first isolated from the *Nmu* gene of C57BL/6J mouse genomic DNA and were validated by sequencing. A positive control vector for subsequent PCR screening assays was also generated, in this case mimicking the DNA configuration of the targeted locus after the homologous recombination event between the short homology arm and the *Nmu* locus. The linearized targeting vector was electroporated into embryonic stem (ES) cells, which were then subjected to positive selection. The resistant ES cell clones were first subjected to a PCR screening over the 3’ short homology arm, using a primer hybridizing within the neomycin cassette and a primer hybridizing downstream of the 3’ homology arm. The optimized PCR protocol was tested in wild-type DNA from C57BL/6J mice as a negative control, and in the positive control vector in the presence of wild-type C57BL/6J genomic DNA. The clones validated by PCR were further analyzed by Southern blot to validate the correct recombination event at the 5’ end of the targeted locus, using a neomycin (internal) probe. No additional randomly integrated copy of the targeting construct was reported in the selected ES clones. The correct 3’ recombination event was also further assessed by Southern blot, in this case using an external probe. The detection of two restriction fragments indicated the heterozygous status of the recombined *Nmu* locus. The PCR- and Southern blot-positive recombined ES clones, further validated by sequencing analysis to ascertain the integrity of the inserted cassettes, were injected into blastocysts to obtain chimeric mice carrying the recombined locus. Highly chimeric males assessed by coat color markers comparison were placed in breeding to ensure germline transmission. The selected chimeric animals were crossed with C57BL/6J Flp deleter mice, in order to excise the neomycin selection cassette and generate heterozygous mice carrying the neo-excised knock-in allele. The Flp line was developed at genOway by pronuclear injection of a CMV-Flp encoding sequence. The Flp deleter line displays ubiquitous expression of Flp recombinase and efficient FRT-flanked sequence recombination. It has been used for several years for neomycin cassette removal. The excision of the neomycin selection cassette prevents unwanted interactions between the cassette and control regions of the targeted locus, thus disrupting the expression of neighboring genes within the locus^47^. Progeny was then genotyped by PCR and the excision event, and the integrity of the targeted region was further verified by sequence analysis on a subset of the PCR-validated animals.

**Figure 1.**
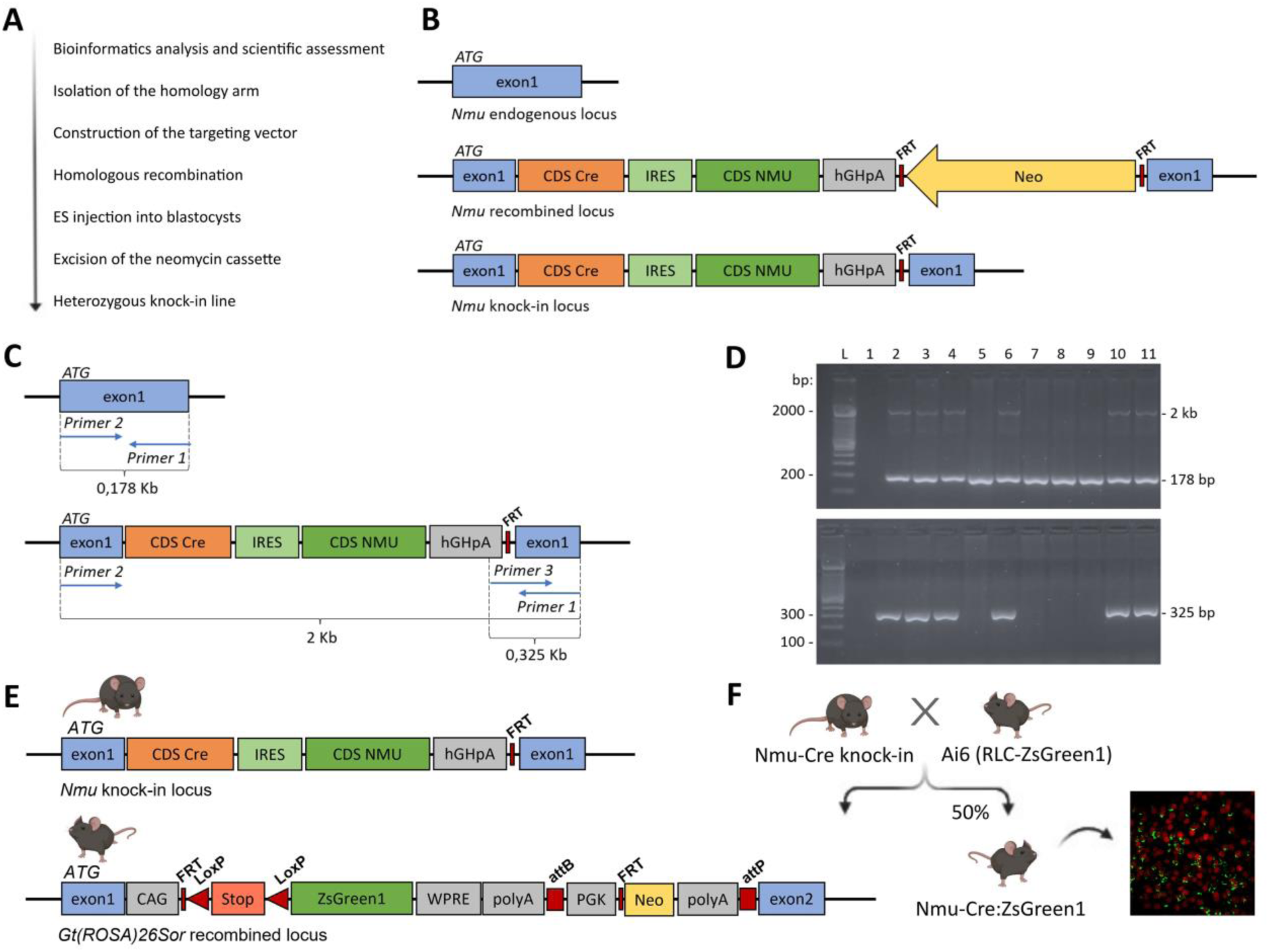
Generation of the Nmu-Cre knock-in mouse model. (A) Step by step progression for the generation of the knock-in line. (B) Schematic representation of the selected *Nmu* gene targeting strategy. The Cre-IRES-*nmu*-hGHpolyA cassette was inserted in frame with exon 1 ATG of the *Nmu* gene. The neomycin selection cassette was excised to generate heterozygous mice carrying the neo-excised *nmu* knock-in allele. (C) Diagram representing *Nmu* endogenous locus (top) and knock-in locus (bottom) with the binding sites of the screening primers. (D) Representative PCR results from genotyped animals. After gel electrophoresis, a distinction could be made between wild-type and heterozygous genotypes. PCR products of 178 bp, 325 bp and 2 kb (lines 2-4, 6, 10 and 11) corresponding to the heterozygous genotype; PCR products of 178 bp (lines 5, 7-9) corresponding to the wild-type genotype. L: ladder; line 1: negative control. (E) Schematic representation of the *Nmu* knock-in locus in Nmu-Cre mice (top) and the *Gt(ROSA)26Sor* recombined locus in Ai6 reporter mice (bottom). (F) Representation of the cross between Nmu-Cre knock-in mice and Ai6 reporter mice. The resulting offspring, Nmu-Cre:ZsGreen1, express the green fluorescent protein ZsGreen1 in all cells expressing Cre recombinase. Diagrams are not depicted to scale. IRES: internal ribosome entry site; CDS: coding sequence; FRT: flippase recognition target sites; Neo: neomycine resistance gene; WPRE: woodchuck hepatitis virus post-transcriptional regulatory element; PGK: phosphoglycerate kinase selection marker; attB/attP: specific recombination sites; LoxP: locus of X(cross)-over in P1.

### 2.3. Genotyping

Mutant mice were offspring of the cross between Nmu-Cre knock-in and C57BL/6J mice or between Nmu-Cre knock-in and Ai6 reporter mice. Genotypes of the offspring were verified by PCR amplification of genomic ear DNA using the commercially available REDExtract-N-Amp™ Tissue PCR Kit (#R4775; Sigma-Aldrich, Germany) and primers designed to identify the presence or absence of the knock-in allele. The following primer couples were used: 5’-GTGACAGGAGAGGAGATGCGGTTGC-3’ (forward primer) and 5’-AGCAAGAGGAGGCGCACAGGA-3’ (reverse primer) to detect the wild-type allele (178 bp); and 5’-GTGACAGGAGAGGAGATGCGGTTGC-3’ (forward primer) and 5’-ACCTTGGCCTCCCAAATTGCTG-3’ (reverse primer) to detect the neo-excised knock-in allele (325 bp) (Figure 1C, D). All primers were obtained from Eurogentec (Belgium). The PCR amplification reaction took place at the following conditions: 94°C for 2 min, 30 cycles of 94<u>°</u>C for 30 s, 65<u>°</u>C for 30 s and 68<u>°</u>C for 5 min, and a final extension step at 68<u>°</u>C for 8 min. PCR reaction products were separated on a 2 % agarose gel electrophoresis and visualized with GelRed (nucleic acid gel stain; #41003, Biotium, USA).

### 2.4. Viral transduction

Viral transduction of NMU-expressing cells was achieved by intracerebral injection of the ready-to-use double-floxed adeno-associated viral (AAV) vector AAV5-EF1a-DIO-RFP (Vector Biosystem, USA), driving Cre-dependent expression of the red fluorescent protein (RFP) in a temporally and regionally defined manner. Hereto, mice were deeply anesthetized in an induction chamber with 2-3 % isoflurane (1000 mg/g, Vetflurane Neurology, Virbac, Belgium) and mounted on a stereotaxic frame. Anaesthesia was maintained during the entire duration of the surgery using 1-2 % isoflurane in 100 % oxygen, delivered via an inhalation cone. Meloxicam (5 mg/kg, Metacam^®^, 5 mg/mL, Boehringer Ingelheim, Germany) was administered subcutaneously to prevent postoperative pain and inflammation. Artificial tear ointment (Duratears^®^, Novartis, Belgium) was applied on the eyes to prevent dehydration. After disinfection of the skin with iso-Betadine^®^ (Mylan, Belgium), an incision was made, the skull was cleared and small holes were drilled in the skull (Volvere Vmax, NSK, Germany) at specific coordinates relative to bregma (Table 1). Next, the AAV vector was bilaterally infused into the target regions at a flow rate of 0,15 μL/min using a 10 μL microsyringe (Hamilton Neuros, USA). The syringe was kept in place for an additional 10 min to limit backflow along the injection track. After injections, the skin was closed using nonabsorbable suture (Ethilon II, 4-0, M-2, Ethicon, USA). At the end of the surgical procedure, mice received 1 mL saline (0,9 % NaCl, Baxter, Belgium) intraperitoneally to prevent dehydration and were placed in a recovery box with heating (ThermaCage^®^, Datesand Group, UK) until fully awake and responsive. After surgery, mice were returned to their home cage to allow further recovery. Mice were single-housed to prevent damage to the suture and sacrificed for *ex vivo* experiments 3 weeks after surgery to ensure proper transduction of the AAV vector and full expression of RFP.

**Table 1.**
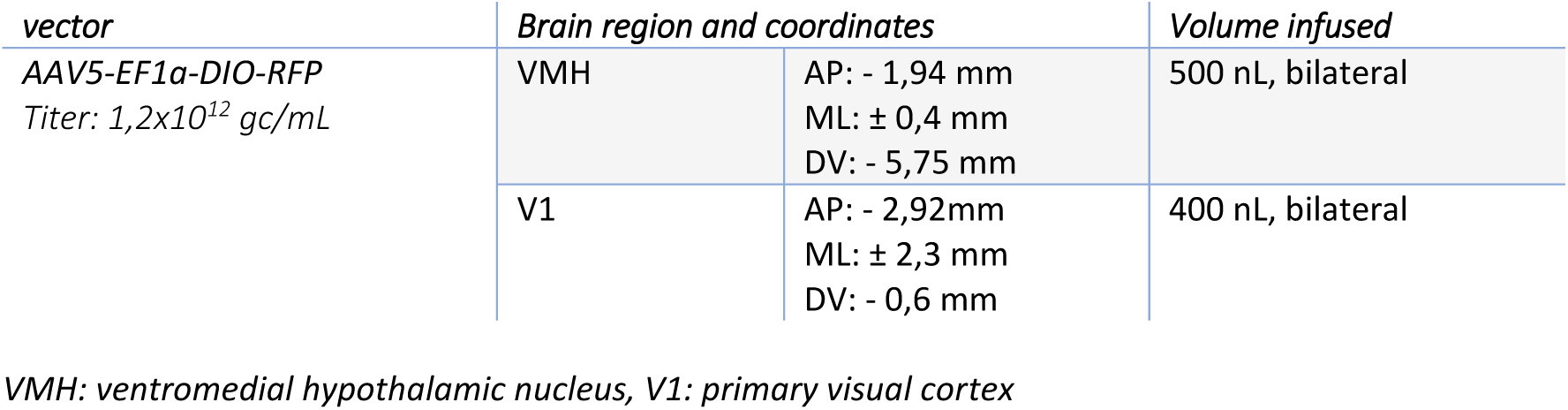
Viral vector infusion and the corresponding coordinates relative to Bregma.

| *vector* | *Brain region and coordinates* |  | *Volume infused* |
| --- | --- | --- | --- |
| AAV5-EF1α-DIO-RFP<br>Titer: $1,2 \times 10^{12}$ gc/mL | VMH | AP: - 1,94 mm<br>ML: ± 0,4 mm<br>DV: - 5,75 mm | 500 nL, bilateral |
| | V1 | AP: - 2,92mm<br>ML: ± 2,3 mm<br>DV: - 0,6 mm | 400 nL, bilateral |

VMH: ventromedial hypothalamic nucleus, V1: primary visual cortex

### 2.5. Tissue preparation for histology

Mice were deeply anesthetized by an intraperitoneal overdose of sodium pentobarbital (250 mg/kg Dolethal^®^, Vetoquinol, France). Immediately after respiratory arrest, transcardial perfusion was performed using phosphate-buffered saline (PBS, pH 7,4, Sigma-Aldrich, Germany) followed by 4 % paraformaldehyde (PFA, VWR International, Belgium) in PBS (pH 7,4) for 5 min at a rate of 10 mL/min. After transcardial perfusion, brains were dissected and postfixated overnight with PFA 4 % and then stored in Tris-buffered saline (TBS) solution (50 mM Tris, pH 7,6, Sigma-Aldrich, Germany) at 4°C. Forty μm coronal sections were sliced using a vibratome (Leica VT1000S, Leica Biosystems, Germany) and stored at −20 °C in an anti-freeze solution (30 % glycerol [Merck Millipore, Germany], 30 % ethylene glycol [VWR International, USA] and 10 % TBS) until further processing. Free-floating sections of specific brain areas were selected and rinsed three times with TBS, each for 10 min. Next, sections were pretreated with TBS containing 0,1 % Triton-X (TBS-T; Sigma-Aldrich, Germany) for 15 min at room temperature, and were then incubated with 25 μg/mL of DAPI (4’,6-diamidino-2-phenylindole dihydrochloride; Cell Signaling Technology, USA) for 5 min. Finally, the slices were washed two times with Tris buffer (TB, 50 mM Tris, pH 7.6, Sigma-Aldrich, Germany), mounted on Superfrost slides (Superfrost plus, VWR international, Belgium) and coverslipped using Dako mounting medium (Agilent, USA). Endogenous ZsGreen1 fluorescence and DAPI staining were evaluated using a fluorescence microscope (Evos FL Auto, ThermoFisher Scientific, USA) and analyzed using Image J (NIH, USA; RRID:SCR_003070).

### 2.6. Immunohistochemistry

Free-floating 40 μm coronal sections were selected and rinsed three times for 10 min with TBS. Next, sections were pretreated with TBS-T containing 10 % donkey serum (Merck Millipore, USA) for 1 h at room temperature under gentle agitation. Next, sections were incubated overnight in primary antibodies (Table 2) at 4 °C. The next day, sections were rinsed three times with TBS-T and incubated with secondary antibodies (Table 3) for 45 min at room temperature and protected from light. DAPI staining and mounting were performed as described in 2.5. Fluorescent labelling was visualized with a fluorescence microscope (Evos FL Auto, ThermoFisher Scientific, USA) and a confocal laser scanning microscope (Zeiss, Axio Observer with LSM 710-6NLO configuration, Zeiss International, Germany), and analyzed using Image J (NIH, USA; RRID:SCR_003070). The specificity of the anti POMC antibody was verified earlier^48^. The specificity of all the other primary antibodies used in this study was determined by Western blot analysis (see supplier’s information, Table 2).

**Table 2.** Primary antibodies.

| *Antibody* | *Class* | *Host species* | *Manufacturer (catalogue nº)* | *Dilution* | *RRID\** |
| --- | --- | --- | --- | --- | --- |
| Anti-Glial Fibrillary Acidic Protein (GFAP) | Polyclonal | Chicken | OriGene, USA (TA309150) | 1:1000 |  |
| Anti-Oligodendrocyte transcription factor 2 (Olig2) | Polyclonal | Rabbit | Merck Millipore, USA (AB9610) | 1:1000 | AB_570666 |
| Anti-Neuronal nuclear protein (NeuN) | Polyclonal | Guinea-pig | Synaptic Systems, Germany (266004) | 1:400 | AB_2619988 |
| Anti-Ionized calcium-binding adaptor molecule 1 (Iba1) | Monoclonal | Rabbit | Thermo Fisher, USA (MA5-27726) | 1:1000 | AB_2735228 |
| Anti-mCherry | Polyclonal | Goat | LifeSpan BioSciences, USA (LS-C204207) | 1:500 | AB_2619713 |
| Anti-Estrogen receptor alpha (ERα) | Polyclonal | Rabbit | Merck Millipore, USA (06-935) | 1:1000 | AB_310305 |
| Anti-Proopiomelanocortin (POMC) | Polyclonal | Rabbit | Phoenix Pharmaceuticals, USA (H029-30) | 1:400 | AB_2307442 |
| Anti-Neuropeptide Y (NPY) | Monoclonal | Mouse | Santa Cruz, USA (sc-133080) | 1:500 | AB_2298657 |

\* RRID: Research Resource Identifier - https://scicrunch.org/resources

**Table 3.** Secondary antibodies.

| *Conjugate* | *Target* | *Host species* | *Catalogue nº* | *Dilution* | *RRID\** |
| --- | --- | --- | --- | --- | --- |
| Cyanine Cy<sup>TM</sup>3 | Chicken | Goat | 103-165-155 | 1:500 | AB_2337386 |
| Cyanine Cy<sup>TM</sup>5 | Rabbit | Donkey | 711-175-152 | 1:400 | AB_2340607 |
| Cyanine Cy<sup>TM</sup>5 | Guinea-pig | Donkey | 706-175-148 | 1:500 | AB_2340462 |
| Cyanine Cy<sup>TM</sup>3 | Rabbit | Donkey | 711-165-152 | 1:400 | AB_2307443 |
| Cyanine Cy<sup>TM</sup>3 | Goat | Donkey | 705-165-147 | 1:400 | AB_2307351 |
| Cyanine Cy<sup>TM</sup>5 | Mouse | Goat | 115-175-146 | 1:400<br>1:800 | AB_2338713 |

All secondary antibodies used were ordered from Jackson ImmunoResearch (USA).
\* RRID: Research Resource Identifier - https://scicrunch.org/resources

### 2.7. Preparation of optically cleared brains

#### 2.7.1. Sample preparation and delipidation

Mice were perfused and their brains collected and postfixed as described above (2.5). Postfixed brains were rinsed with PBS at least 2 times for 2 h at room temperature. Brains were then delipidated with clear unobstructed brain imaging cocktail and computational analysis (CUBIC)-L clearing solution (10 wt% N-buthyldiethanolamine [[#471240, Sigma-Aldrich, Germany] and 10 wt% Triton X-100 [Sigma-Aldrich, Germany] in Milli-Q water)^49,50^. To do so, brains were first immersed in 1:1 Milli-Q water-diluted CUBIC-L with gentle shaking (75 rpm) at 37°C overnight. Next, the samples were incubated in complete CUBIC-L for 10 to 12 days, with gentle shaking (75 rpm) at 37°C. CUBIC-L was refreshed every 2-3 days. Brains were then washed three times for 2 h with PBS at room temperature.

#### 2.7.2. Refractive index matching with CUBIC-R+(M) and agarose gel embedding

Refractive index matching was achieved by incubating the cleared brains in CUBIC-R+(M) (45 wt% 2,3-dimethyl-1-phenyl-5-pyrazole [Antipyrine; #10784, Sigma-Aldrich, Germany], 30 wt% N-methylnicotinamide [#M0374, TCI, Japan] and 0.5 wt% N-buthyldiethanolamine [#471240, Sigma-Aldrich, Germany], pH 9,6-9,8 in Milli-Q water)^50,51^. To do so, the samples were first immersed in 1:1 Milli-Q water-diluted CUBIC-R+(M) with gentle shaking (75 rpm) at room temperature overnight. The next day, the solution was replaced by complete CUBIC-R+(M) and samples were further incubated for 24 h. Gel embedding was based on Matsumoto and colleagues^52^. Briefly, agarose (2 wt/v%) was thoroughly diffused in fresh CUBIC-R+(M) (adjusted pH 8,5-9) and the mixture was then incubated in a hot water bath at 60 °C during the whole embedding protocol. A bottom gel layer was formed in a custom-made mold and was incubated at room temperature for 15 min. Then, the mixture was poured into a 15 mL falcon tube, the sample was added and immediately decanted into the gel mold, positioning the sample in the center and the desired orientation. Bubbles, if present, were carefully removed with a pipette tip and the gel was incubated at room temperature for another 15 min. To prepare the top layer, agarose-CUBIC-R+(M) solution was decanted into the gel mold until the surface protruded slightly. Bubbles were carefully removed, and the embedded sample was incubated at room temperature for at least 4 h.

#### 2.7.3. Refractive index matching with ethyl cinnamate

The samples were gently removed from the mold and incubated for 30 min in fresh CUBIC-R+(M) (adjusted pH 8,5-9). Then, the brains were dehydrated in increasing concentrations of 1-propanol pH 9 (1-propanol in CUBIC-R+; 25, 50, 75, 2 × 100 %, 30 min each step, and a final overnight step at 100 %). The next day, dehydrated samples were incubated in increasing concentrations of ethyl cinnamate (ECi, #112372, Sigma-Aldrich, Germany; ECi in 1-propanol pH 9; 25, 50, 75, 2 × 100 %, 45 min each step)^53–55^ and were kept in 100 % ECi until image acquisition in a Light-sheet Fluorescence Microscope (LSFM).

#### 2.7.4. LSFM image acquisition and processing

Image acquisition was done in ECi using a Lavision Ultramicroscope II (Lavision Biotec, Germany) equipped with an Olympus MVPLAPO 2× (NA 0.50) objective lens and DBE-corrected LV OM DCC20 dipping cap. Images were recorded with a Neo sCMOS camera (Andor), using a 2x objective and a digital zoom of 0,63x. Samples were optically sectioned using a z-step size of 10 μm, resulting in 5,08 μm × 5,08 μm × 10 μm voxel size (16 bit per pixel). A 488 nm and 561 nm laser were used in combination with a 525/50 nm and 620/60 nm emission filter, respectively. Sagittal optical sections were acquired in a mosaic of four tiles using the left and right light sheets separately, respectively for the left and right part of the mosaic. The resulting image sets were then imported into Arivis Vision 4D software (Zeiss, Germany; RRID SCR_018000) for stitching and acquisition of movies, images and 3D renderings.

### 2.8. RNA *in situ* hybridization

#### 2.8.1. Chromogenic detection

Adult Nmu-Cre mice were perfused as described in 2.5, using 10 % neutral buffered formalin (NBF) as a fixative. After transcardial perfusion, brains were dissected and postfixated for 24 h with 10 % NBF at room temperature in a rotating shaker, followed by standard paraffin embedding using the automatic tissue processor Leica TP 1020 (Leica Microsystems, Germany). Brains were sectioned with a microtome at 5 μm thickness. The RNAscope^®^ 2.5 HD Assay-RED (#322350, Advanced Cell Diagnostics, USA) was used according to the manufacturer’s instructions using adapted pretreatment conditions (15 min Target Retrieval and 5 min Protease Plus). Probes used were *Bacillus subtilis* dihydrodipicolinate reductase (dapB, #310043), *Mus musculus* peptidyl-prolyl isomerase B (Ppib, #313911) and *Mus musculus* Nmu (#446831). Slides were counterstained for hematoxylin and mounted with VectaMount^®^ mounting medium (Vector Laboratories, USA). Images were acquired with the brightfield microscope Olympus BX61 (Japan) at 20x and 40x magnification.

#### 2.8.2. Fluorescent multiplex detection

Adult Nmu-Cre mice were sacrificed by cervical dislocation, and brains were removed, placed immediately on dry ice for 5 min and stored at −80°C. Brains were sectioned at −17°C with a cryostat at 14 μm thickness. Slices were collected onto Superfrost Plus slides (ThermoFisher, USA). Probes for *Nmu* (#446831-C3), *Slc32a1* (#319191-C2) and *Slc17a6* (#319179-C1) were used with the RNAscope^®^ Fluorescent Multiplex labeling kit (#320850; Advanced Cell Diagnostics) according to the manufacturer’s recommendations. Slides were mounted with ProLong Diamond Antifade mountant (#P36961, Invitrogen, USA). Single-molecule fluorescence of labeled cells were captured using sequential laser scanning confocal microscopy (Leica SP8, Germany) and analyzed using Image J (NIH, USA; RRID:SCR_003070).

### 2.9. Quantitative reverse-transcription polymerase chain reaction (RT-qPCR)

RNA extraction, reverse transcription and quantitative PCR were performed as previously reported^56^ with slight modifications. Mice were sacrificed by cervical dislocation and brains were rapidly removed from the skull. The hypothalamus and hippocampus were dissected and snap-frozen in 2-methylbutane (J.T. Baker, Poland) and stored at −80 °C. Extraction and purification of total RNA was performed using the RNeasy Mini Kit (#74104, Qiagen, Germany) according to the manufacturer’s instructions. A Nanodrop spectrophotometer (Thermo Scientific, USA) was used to check relative RNA concentration and purity, measuring both 260/280 nm (~2.0) and 260/230 nm (~2.0-2.2) ratios. Following extraction, complementary DNA (cDNA) synthesis (iScript cDNA synthesis kit, Biorad,, Belgium) and cDNA purification (Genelute PCR clean-up kit, Sigma-Aldrich) were carried out, according to the manufacturer’s instructions. qPCR was performed using SYBR™ Green PCR master mix (#4309155, Applied Biosystems™, ThermoFisher, USA). Specific primers for the gene of interest *Nmu* (NCBI reference sequence: NM_019515.1; forward 5’-ATCTGTTGCACTGAGGGAGC-3’; reverse 5’-TCATGCAGTTGAGGGACGAG-3’) and the reference genes beta-2 microglobulin (*B2m*, NCBI reference sequence: NM_009735-3; forward 5’-TTCTGGTGCTTGTCTCACTGA-3’; reverse 5’-CAGTATGTTCGGCTTCCCATTC-3’) and hypoxanthine guanine phosphorybosyl transferase 1 (*Hprt1*, NCBI reference sequence: NM_013556.3; forward 5’-CAAACTTTGCTTTCCCTGGT-3’; reverse 5’-TCTGGCCTGTATCCAACACTTC-3’) were used. The resulting amplicon sizes were 186 bp, 104 bp and 101 bp, respectively. All primers were obtained from Eurogentec (Belgium). All qPCR samples were loaded in duplicate. Amplifications were performed using QuantStudio™ 3 Real-time PCR System (Applied Biosystems™, ThermoFisher, USA). qBase+ software (Biogazelle, Belgium, RRID SCR_003370) was used to identify stably expressed reference genes^56^, and were subsequently used for gene normalization. Calculation of relative gene expression was done using the Pfaffl method^57^, taking into account primer efficiencies. Sample sizes were calculated a priori using G*Power (RRID:SCR_013726) and based on standard deviations from previous findings within the research group^56^.

### 2.10. Statistical analysis

Graphical representations and statistical analyses were performed using GraphPad Prism 8 software (GraphPad Software, Inc., USA, RRID SCR_002798). Data are expressed in columns with dots representing individual values, with designation of mean with 95% confidence interval. Data were analyzed using t-tests (unpaired, two-tailed). Significance threshold was set at alpha = 0.05.

## 3. Results

### 3.1. Generation of a Nmu-Cre knock-in mouse model

The Nmu-Cre knock-in mouse was created by the insertion of a Cre-IRES-*Nmu* cassette in frame with the initiation codon of exon 1 of the *Nmu* gene (Figure 1A, B). This targeting strategy with an IRES site generates one single mRNA containing both coding sequences, thus mediating bicistronic translation. As a result, Cre and NMU will be expressed as discrete proteins, being Cre expression regulated by the endogenous *Nmu* promoter. All Nmu-Cre heterozygous knock-in mice obtained were identified by genotyping (Figure 1C, D). Mice were healthy, viable and fertile. The animals displayed no distinct abnormalities or noticeable developmental defects, based on visual assessment of physical appearance, body condition, unprovoked behavior and brain morphology. However, continuous monitoring is recommended to detect any potential minor phenotype.

### 3.2. Neuroanatomical characterization of ZsGreen1-expressing cells in brain coronal slices from Nmu-Cre:ZsGreen1 mice

Heterozygous Nmu-Cre mice were crossed with homozygous Ai6(RCL-ZsGreen) reporter mice to visualize and characterize NMU-containing cells in the Nmu-Cre mouse model^41^ (Figure 1E, F, Suppl. Figure 1). Offspring of this cross were identified by genotyping (Figure 1C, D). Nmu-Cre:ZsGreen1 mice were found to be healthy and viable. To assess the possible spontaneous recombination between LoxP sites and define ZsGreen1 fluorescence background levels, we analyzed coronal slices obtained from male and female Ai6 homozygous mice. Minimal and sparse ZsGreen1 signal was observed, consistent with a very low spontaneous recombination rate in the absence of Cre recombinase (Suppl. Figure 2A). Next, coronal slices of both female (Figure 2) and male (Suppl. figure 2B-C) Nmu-Cre:ZsGreen1 mice were systemically analyzed by fluorescence microscopy, examining 25 to 35 coronal slices per mouse brain at similar antero-posterior levels. The expression of ZsGreen1 appeared to be stable across litters and consistent between sexes. ZsGreen1 was distributed throughout the cortex, and in subcortical regions comprising the hypothalamus, amygdala, midbrain and brainstem (Figure 2A-B, Suppl. Figures 1-2). As previously described, ZsGreen1 fluorescence was mainly detected in the cell bodies, and displayed a punctate labeling pattern^41^. Remarkably, ZsGreen1 was photostable, unaffected by fixation, and was readily detected without further amplification^58^. A qualitative gradation was used to describe mainly absent (−), low (+), moderate (++) and high (+++) ZsGreen1 levels (Table 4). The nomenclature used in the text, figures and tables follows the mouse brain atlas of Paxinos and Franklin^59^. All the brain regions examined for endogenous expression of ZsGreen1 are listed in Table 4 (see Table 5 for explanation of other abbreviations). ZsGreen1-expressing cells were observed throughout the cortex, from anterior to posterior levels. While only a few marked cell bodies were detected in the medial prefrontal cortex (mPFC), moderate ZsGreen1 expression levels were found in motor and somatosensory areas, located mainly in layers II/III, IV and V. Remarkably, a high and dense signal was observed in the visual cortex, comprising both primary (V1) and secondary (V2) visual areas. Very low ZsGreen1 expression was found in the nucleus accumbens (NAc), located sparsely in the shell, and dorsally, in the lateral septum (LS), while it was mainly absent in striatal, basal forebrain and hippocampal regions. In contrast, moderate to high ZsGreen1 expression levels were detected in the bed nucleus of stria terminalis (BNST). Similarly, moderate ZsGreen1 expression was clearly observed in a few thalamic nuclei surrounding the paraventricular thalamic nucleus (PV), in the medial amygdaloid nucleus (MeA) and in several nuclei within the hypothalamus. The SCh, VMH and the mammillary body (MM), showed the most robust and dense fluorescent expression. ZsGreen1-marked cells were also observed within the midbrain, especially throughout the periaqueductal gray (PAG). Ventral to the aqueduct, neurons within the Edinger-Westphal nucleus (EW) showed moderate expression, which was also detected more ventrally, in the anterior portion of the rostral linear nucleus of the raphe (RLi) and the parabrachial pigmented nucleus (PBP) of the ventral tegmental area (VTA). Low to moderate expression of ZsGreen1 was found in regions within the medulla, such as the parvicellular reticular nucleus, alpha part (PCRtA).

**Figure 2.**
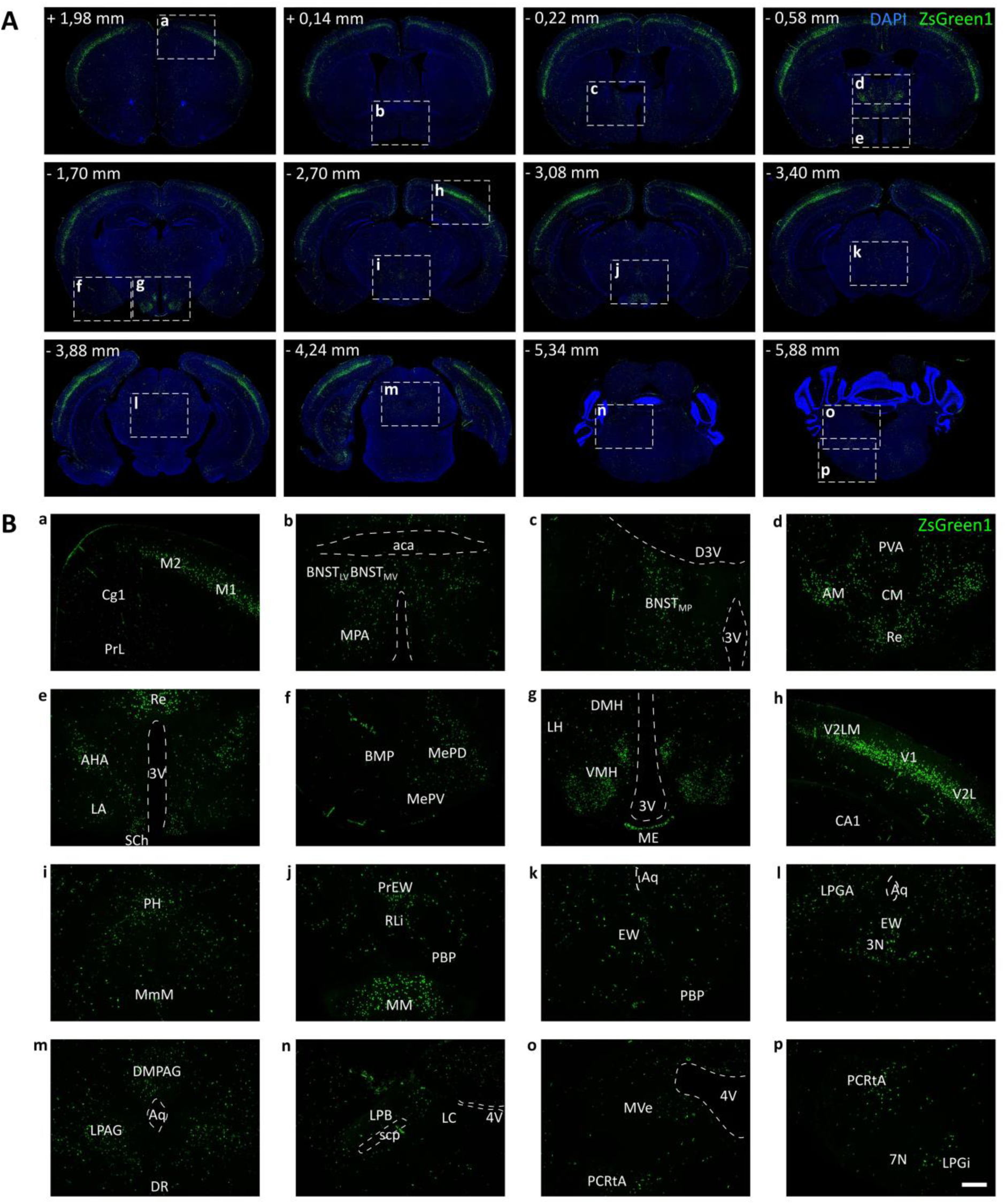
Neuroanatomical distribution of ZsGreen1 expression in coronal brain slices. (A-B) Distribution of ZsGreen1-expressing cells in coronal slices from Nmu-Cre:ZsGreen1 mice (representative images from 1 female; n = 4 females were analyzed). The qualitative analysis was performed on 25-35 coronal slices per mouse brain by two independent observers. DAPI nuclear staining (blue), ZsGreen1 (green). Areas of higher magnification are indicated by a dashed white box. Images were acquired with a fluorescence microscope at 4x (A) and 10x (B) magnification. Numbers denote neuroanatomical coordinates relative to bregma. Scale bar: 300 μm. DAPI: 4’,6-diamidino-2-phenylindole dihydrochloride. Anatomical abbreviations are listed in Table 5.

**Table 4.**
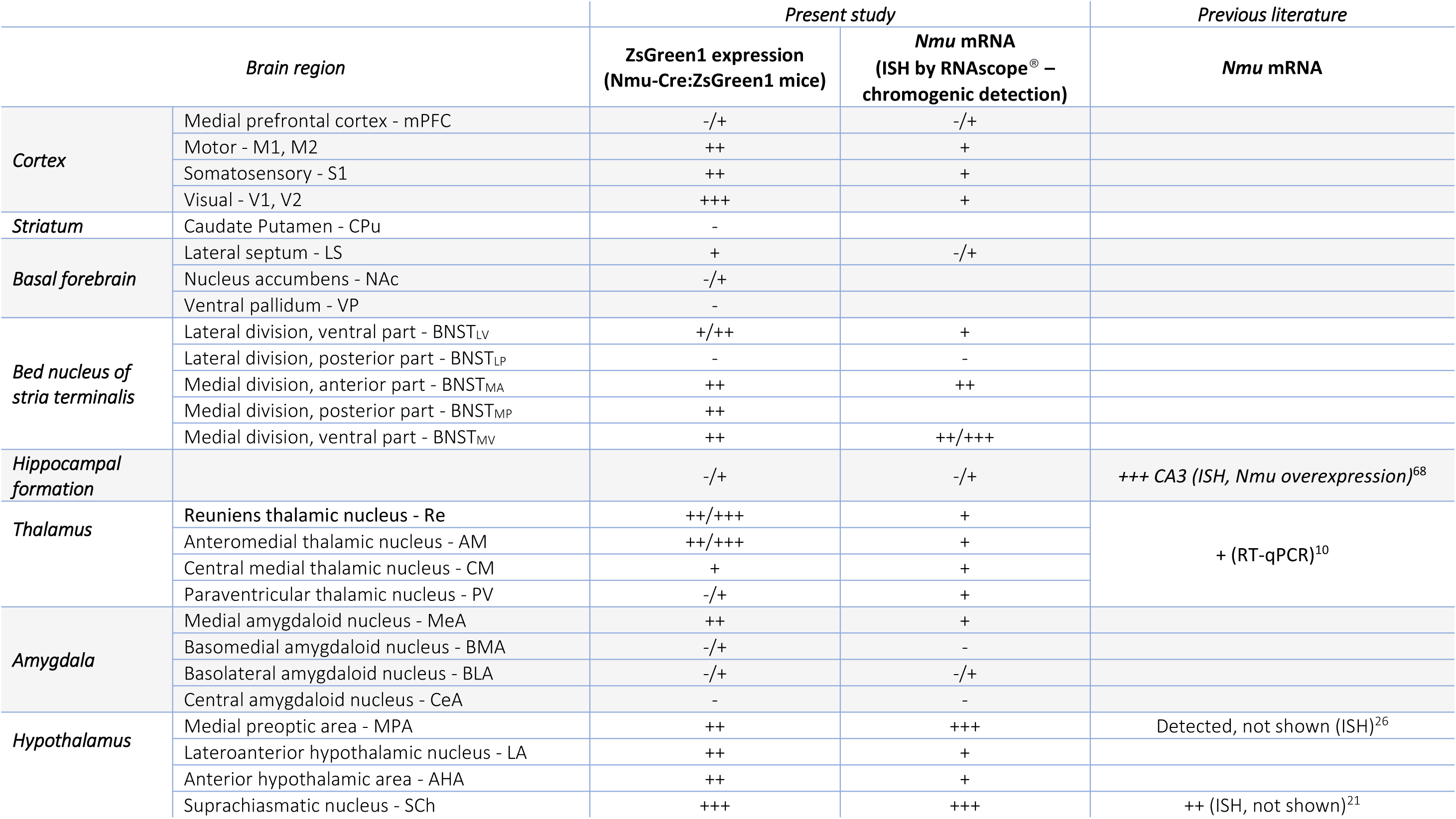
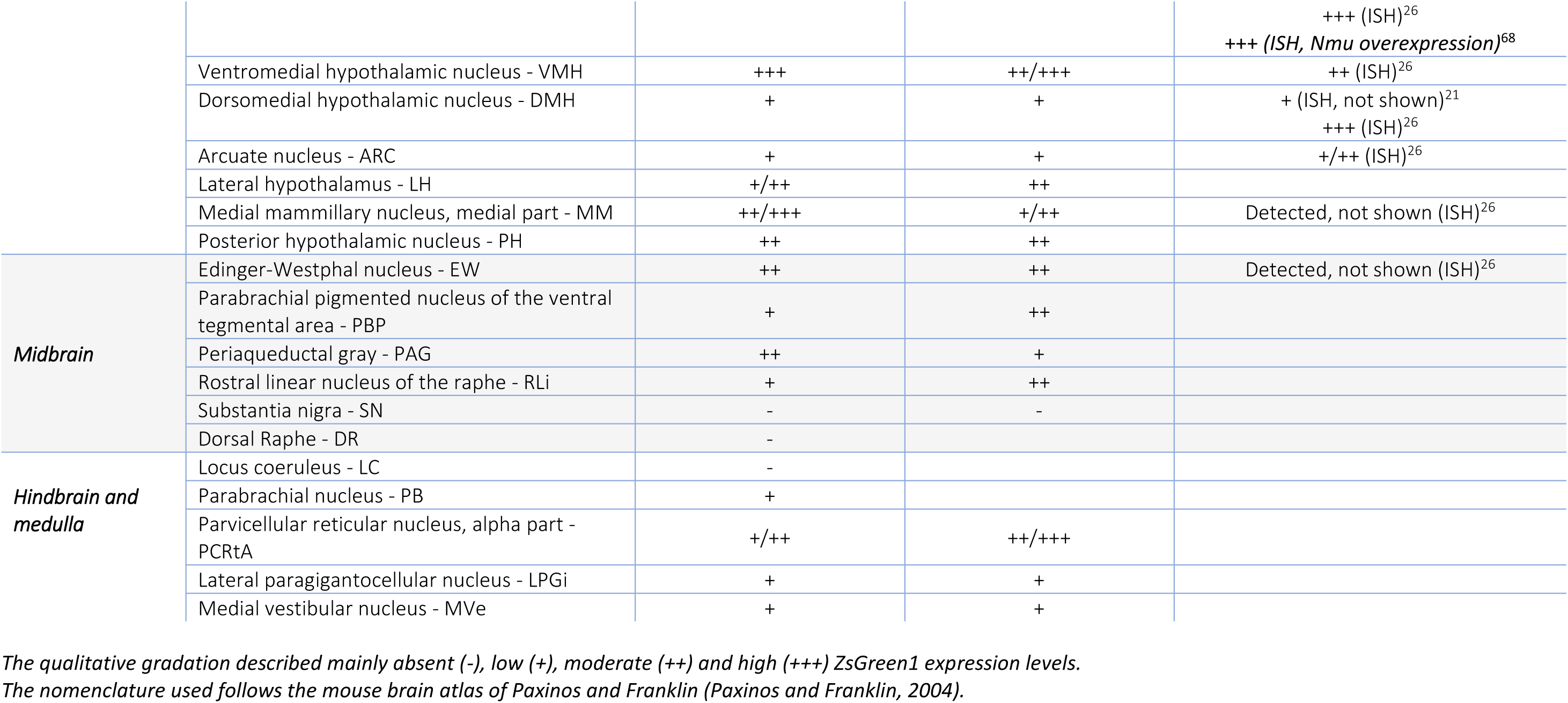
Qualitative analysis of ZsGreen1 fluorescence levels.

**Table 5.**
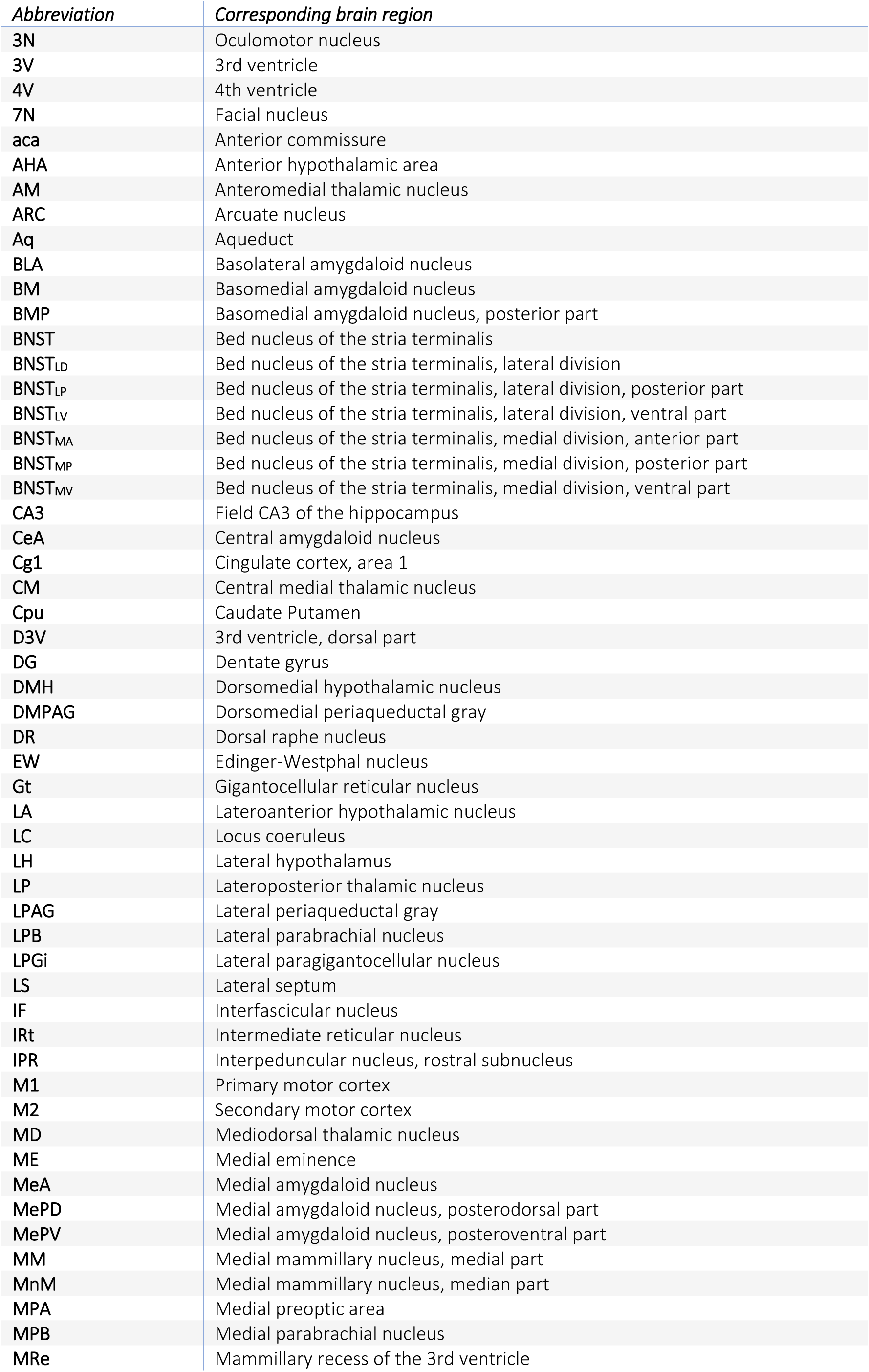
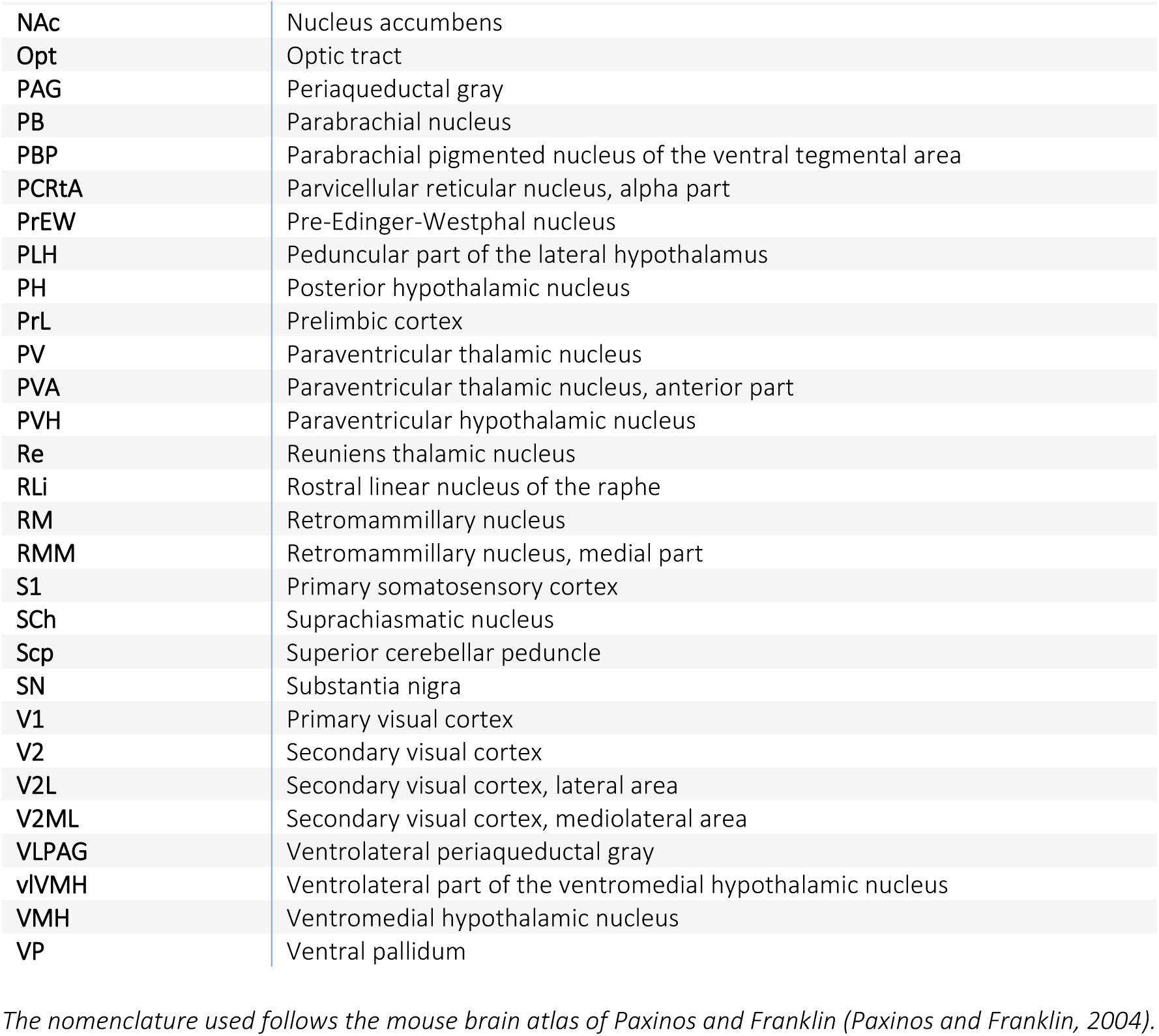
List of brain regions and its abbreviations.

### 3.3. Neuroanatomical characterization of ZsGreen1-expressing cells in cleared mouse brains

To provide a complete picture and a more comprehensive description of ZsGreen1 expression in Nmu-Cre:ZsGreen1 mice, we optimized a new protocol for optimal delipidation, decolouring and refractive index matching based on a combination of existing clearing and RI matching reagents (Figure 3A). Delipidation and decolouring were performed using CUBIC-L^49,50^. CUBIC-L induced limited swelling, which was minimized further during the first RI matching step with CUBIC-R+(M)^50,51^. After dehydration with pH-adjusted 1-propanol series, a last step using the organic solvent ECi was performed, taking advantage of the complementary RI matching effect of both CUBIC-R and ECi^54^. The combination of 1-propanol pH 9 and ECi preserved endogenous fluorescence^53–55^. In the optically cleared tissue, we performed volumetric imaging of ZsGreen1 signal using LSFM. This revealed a continuous population of ZsGreen1-expressing cells extending from the ventral forebrain to the caudal midbrain, comprising numerous regions such as the ST, medial preoptic area (MPA), SCh, VMH, lateral hypothalamus (LH), MeA, posterior hypothalamus (PH), MM, EW, VTA and PAG (Figure 3B, C, Video 1). ZsGreen1 expression extends beyond the midbrain, through the pons and the medulla (Figure 3B, C, Video 1). Therefore, this protocol provides optimal clearing of mouse brain while successfully preserving the endogenous ZsGreen1 fluorescence. The observed distribution pattern was consistent with discrete coronal slices.

**Figure 3.**
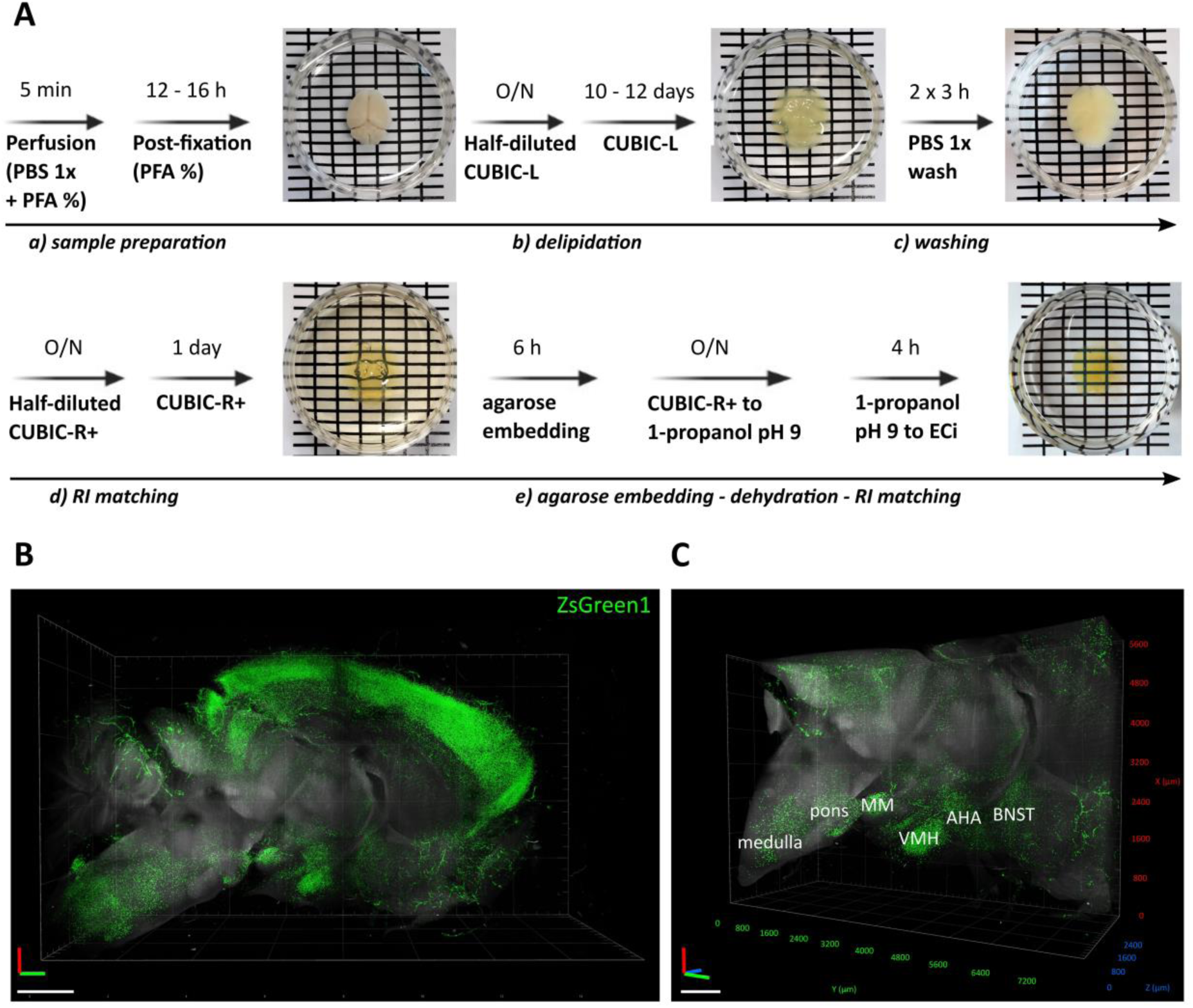
Neuroanatomical distribution of ZsGreen1 expression in cleared mouse brains. (A) Clearing and decoloring protocol using CUBIC-L, and refractive index matching protocol using the complementary effect of both CUBIC-R and ECi. After perfusion, postfixed brains (*a*) were immersed in half-diluted CUBIC-L clearing solution overnight. Lipid removal (*b*) was achieved by treating the brain with complete CUBIC-L for 10 to 12 days, with gentle shaking (75 rpm) at 37°C. The brains were washed in PBS (*c*) and then immersed in half-diluted CUBIC-R+(M) O/N and changed to complete CUBIC-R+(M) for a further 24h (*d*). After the first refractive index matching step, cleared brains were embedded in agarose and final matching was achieved by dehydrating the samples in increasing concentrations of 1-propanol pH 9, followed by increasing concentrations of ECi (*e*). Samples were stored in fresh ECi at room temperature until imaging. (B) Overview 3D volume rendering of half brain from a Nmu-Cre:ZsGreen1 mouse and (C) XYZ-plane clipping of B. Representative images from 1 mouse brain; n = 3 brains were cleared and analyzed. Objective 2x, digital zoom 0,63. 488 nm and 561 nm lasers were used to reveal ZsGreen1 endogenous fluorescence (green) and background signal (gray), respectively. Scale bar: 1 mm. PBS: phosphate-buffered saline; PFA: paraformaldehyde; CUBIC: clear unobstracted brain imaging cocktails and computational analysis; ECi: ethyl cinnamate; O/N: overnight. Anatomical abbreviations are listed in Table 5.

### 3.4. Transcriptional validation of the Nmu-Cre knock-in mouse model

The ZsGreen1 distribution pattern observed in our study was more widespread than anticipated based on literature, although it is comparable with the distribution observed in the Nmu-Cre BAC transgenic mice^42^. It is known that transient developmental expression of Cre can contribute residual fluorescent expression in cells where the *Nmu* promoter is no longer active in adult life^40^. We therefore carried out transcriptional validation experiments.

#### RT-qPCR

The design of the Nmu-Cre knock-in mouse model was based on the absence of regulatory elements that may be impacted by the insertion of the Cre-IRES-*Nmu*-hGHpolyA recombination cassette in exon 1 of the *Nmu* gene. To assess whether the Nmu-Cre knock-in mouse model expressed *Nmu* and did so at similar levels to those observed in wild-type mice, RT-qPCR analyses were carried out. We selected the hypothalamus and hippocampus to assess *Nmu* transcript levels between wild-type and Nmu-Cre knock-in mice and test for regional differences, as described in literature and consistently in Nmu-Cre:ZsGreen1 mice. RT-qPCR results, normalized to the expression of the reference genes *B2m* and *Hprt1*, showed similar relative *Nmu* mRNA expression levels in the hypothalamus of both Nmu-Cre and wild-type mice, indicating that *Nmu* mRNA is indeed expressed in the Nmu-Cre mouse and this expression is not significantly dysregulated in the knock-in mouse model (Figure 4A). To further examine whether ZsGreen1 signal represented the endogenous NMU expression pattern, relative *Nmu* mRNA expression levels were compared between hypothalamus and hippocampus. The results did indeed show significantly higher levels in hypothalamus compared to those found in the hippocampus (Figure 4B).

**Figure 4.**
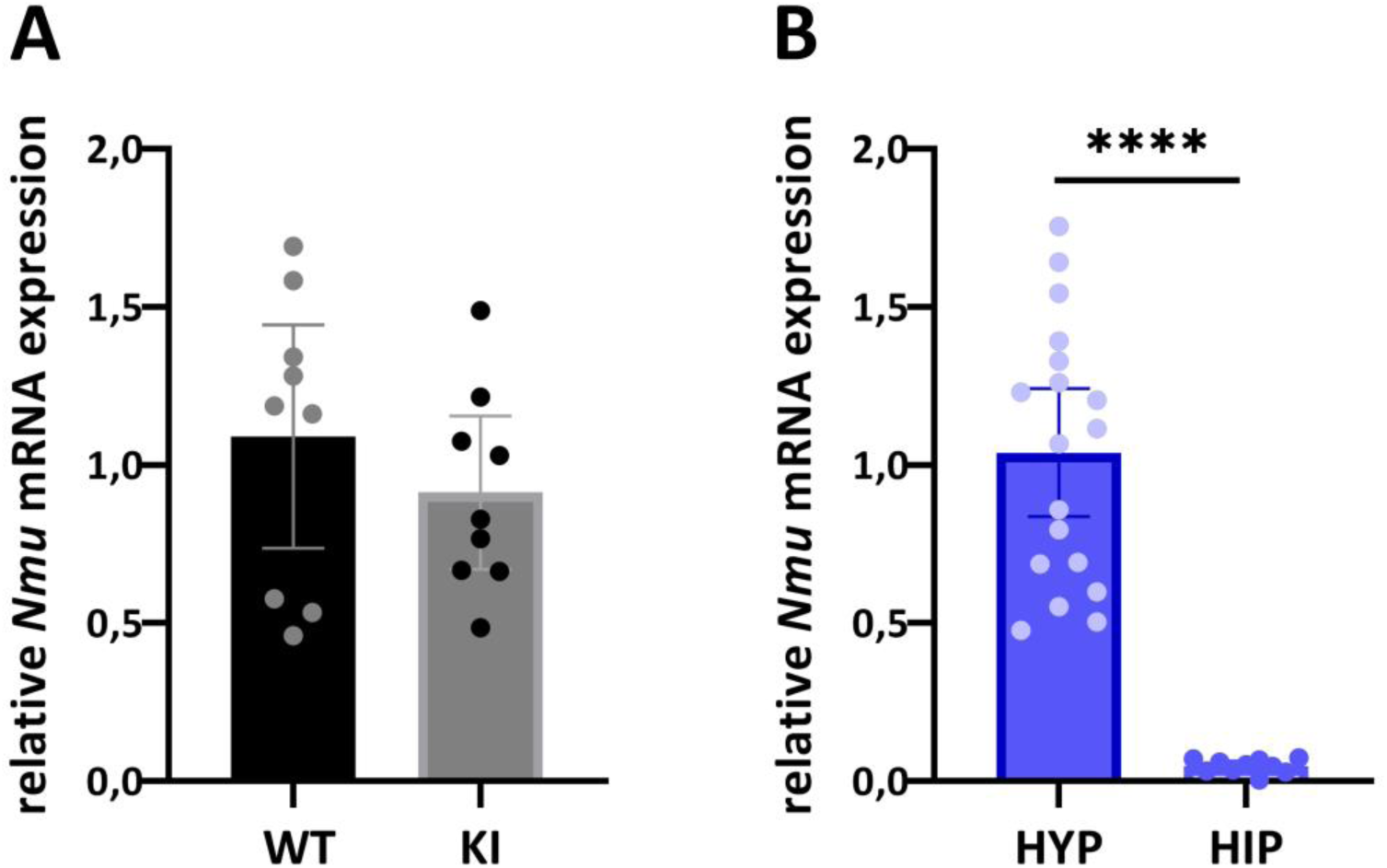
Quantification of the expression of *Nmu* mRNA by using quantitative reverse-transcription polymerase chain reaction (RT-qPCR). (A) Relative *Nmu* mRNA expression in the hypothalamus of wild-type (n = 9) and Nmu-Cre knock-in mice (n = 9). (B) Comparison of *Nmu* mRNA expression in the hypothalamus (n = 18) and the hippocampus (n = 10), considering results obtained from both wild-type and Nmu-Cre mice. The experiment was replicated two times. Data were analyzed with the Pfaffl method and normalized to the reference genes *B2m* and *Hprt1*. Dots represent individual data points; representation of the mean with a 95% confidence interval. Data was analyzed by student’s t-test; ****p<0,0001. WT: wild-type; KI: knock-in; HYP: hypothalamus; HIP: hippocampus.

#### In situ hybridization

To further assess whether ZsGreen1-expressing cells faithfully represented endogenous NMU expression pattern, *Nmu* mRNA expression was analyzed by RNA ISH using RNAscope^®^ technology, a method that provides higher sensitivity and specificity for the detection of low-abundant transcripts than classical ISH techniques^34^. Indeed, while we failed to detect *Nmu* transcript using classical FISH, we did obtain reliable results using RNAscope^®^. Initial attempts to detect *Nmu* mRNA using fluorescent probes and its overlap with ZsGreen1 signal in brain coronal slices from Nmu-Cre:ZsGreen1 mice were unsuccessful. The RNAscope^®^ *in situ* signal detected was unconsistent between mice, and sample processing during the ISH protocol resulted in an almost complete loss of ZsGreen1 fluorescent signal, which prevented us from verifying whether the RNAscope^®^ signal overlapped with the green fluorescent signal. In addition, the lack of suitable anti-ZsGreen1 antibodies hampered the detection of ZsGreen1 after the ISH procedure. Further attempts using RNAscope^®^ and chromogenic detection in Nmu-Cre mice showed consistent results (Figure 5). Each punctate dot signal represents a single *Nmu* mRNA molecule while clusters result from multiple mRNA molecules, visualized in red in contrast with nuclei stained with hematoxylin. Overall, the ISH results in Nmu-Cre mice are largely in agreement with the ZsGreen1 results found in Nmu-Cre:ZsGreen1 mice (Table 4). However, the expression of *Nmu* mRNA was found to be generally lower and more restricted. In line with the ZsGreen1 data, moderate to high *Nmu* mRNA signal was found in several divisions of the BNST, as well as in hypothalamic, midbrain and hindbrain nuclei. The most robust transcript signal was found in the SCh, MPA and VMH. In contrast, very low *Nmu* mRNA expression was observed through the cortex, amygdala and thalamus, where strong ZsGreen1 signal was described. Notably, only a few cells expressing very low amounts of transcript were detected in visual areas, where a moderate and consistent ZsGreen1 signal was observed.

**Figure 5.**
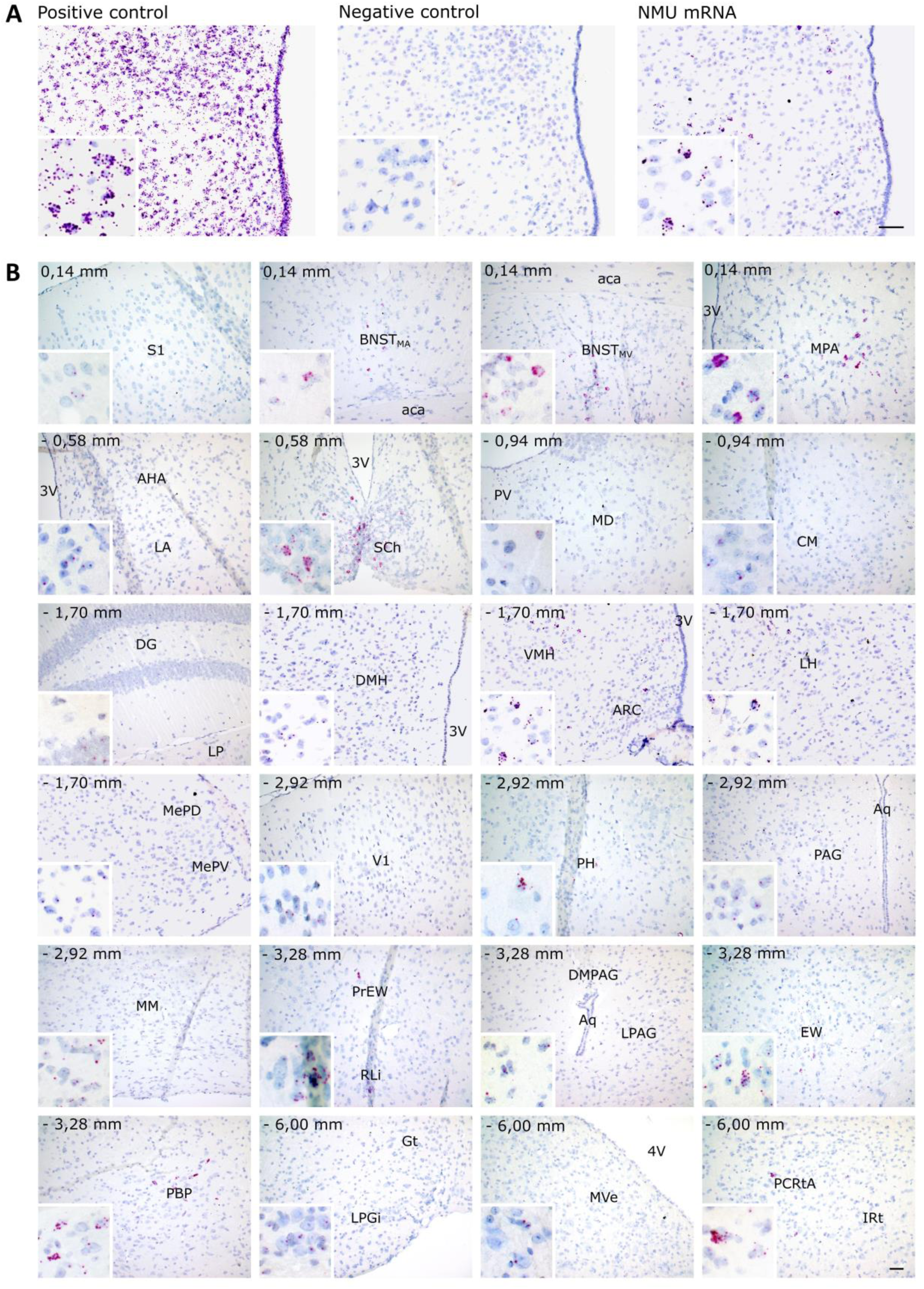
*Nmu* mRNA expression pattern analysis in Nmu-Cre knock-in mice by *in situ* hybridization (RNAscope^®^). (A) Representative images of a positive control (RNAscope^®^ probe against the *Mus musculus* peptidyl-prolyl isomerase B [Ppib]; left), negative control (RNAscope^®^ probe against the *Bacillus subtilis* dihydrodipicolinate reductase [dapB]; middle) and *Nmu* mRNA (right). Hematoxylin nuclear staining (blue); *Nmu* mRNA signal (red). (B) *Nmu* mRNA expression pattern in coronal sections from Nmu-Cre mice. Areas of higher magnification are outlined in white (left-bottom). Representative images from 1 mouse brain; n = 2 brains were analyzed. Images were acquired with the brightfield microscope Olympus BX61 at 20x and 40x magnification. Scale bar: 100 μm. Numbers denote neuroanatomical coordinates relative to Bregma. Anatomical abbreviations are listed in Table 5.

### 3.5. Identification of the cell type expressing NMU

Nmu-Cre:ZsGreen1 mice were used to identify the molecular identity of NMU-expressing cells. Immunostainings using antibodies against glial fibrillary acidic protein (GFAP), ionized calcium-binding adaptor molecule 1 (Iba1), oligodendrocyte transcription factor 2 (Olig2) and neuronal nuclear protein (NeuN) were used as markers for astrocytes, microglia, oligodendrocytes and neurons, respectively. Analysis of colocalization between these specific markers and the fluorescent protein ZsGreen1 were performed in the VMH (Figure 6A) and visual cortex (Figure 7A) of coronal slices using confocal microscopy. As hypothesized, ZsGreen1-expressing cells showed a general overlap with the neuronal marker NeuN but not with the glial markers GFAP, Iba1 and Olig2. These results confirm that NMU-expressing cells are mainly neurons.

**Figure 6.**
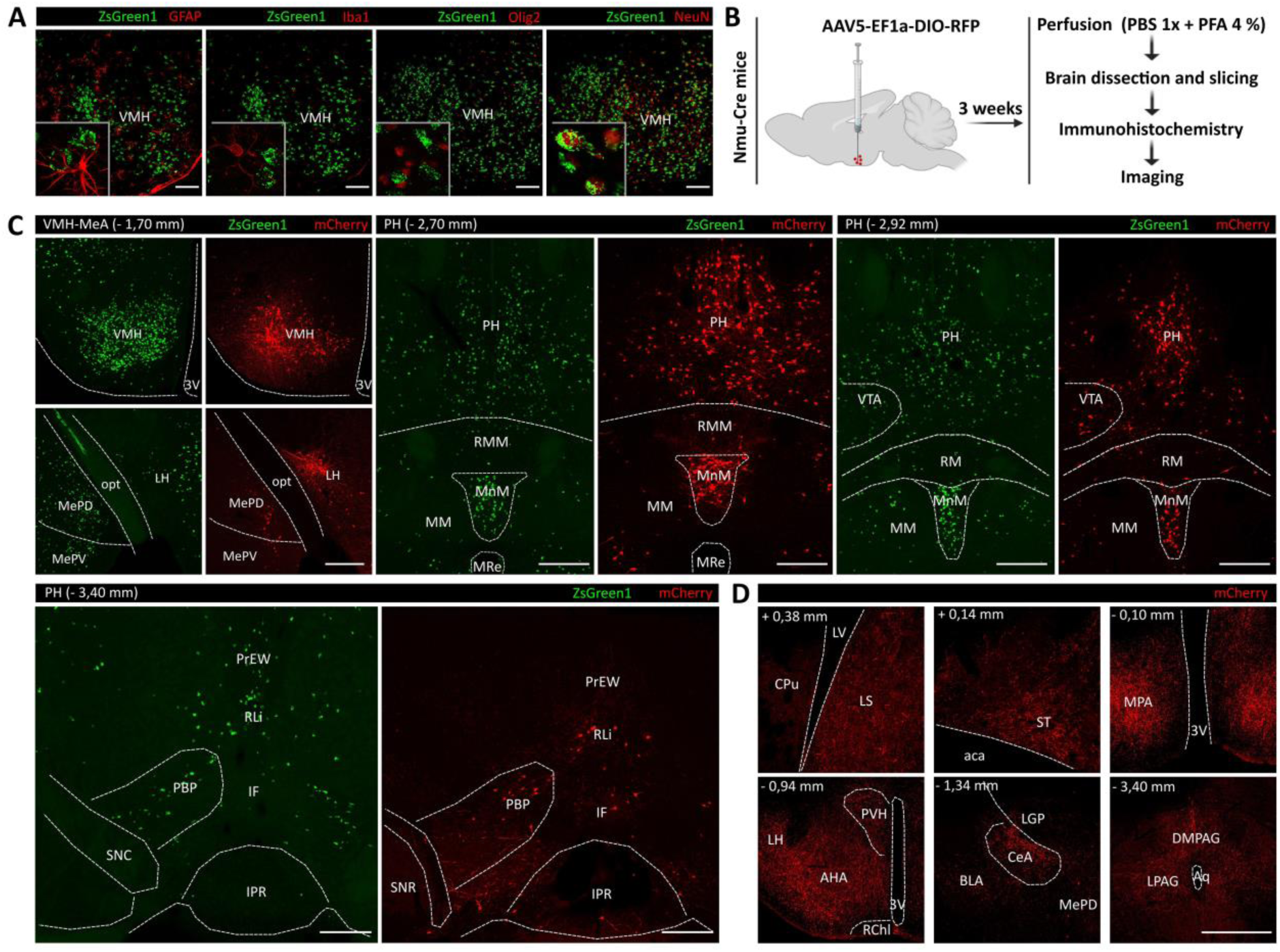
Characterization of NMU neurons in the ventromedial hypothalamic nucleus (VMH) using immunohistochemistry and a viral tracer. (A) Immunohistochemical analysis of brain coronal slices from Nmu-Cre:ZsGreen1 mice at the level of the VMH. In green, ZsGreen1 fluorescence; in red, GFAP and Olig2 (secondary antibodies coupled to Cy3), NeuN and Iba1 (secondary antibodies coupled to Cy5). Representative images from 1 mouse brain; n = 3 brains were analyzed. Images were acquired with a confocal microscope with a 20x objective lens. Areas of higher magnification are outlined in gray (left-bottom). Scale bar: 100 μm. GFAP: glial fibrillary acidic protein; Iba1: ionized calcium-binding adaptor molecule 1; Olig2: oligodendrocyte transcription factor 2; NeuN: neuronal nuclear protein. (B) Schematic representation of the intracerebral injection of the viral tracer AAV5-EF1a-DIO-RFP into the VMH of Nmu-Cre mice. Three weeks after surgery, mice were transcardially perfused, and brains were processed for immunohistochemical amplification of the RFP fluorescent signal with an anti-mCherry primary antibody and a Cy3-conjugated secondary antibody (red). (C) Representative images from coronal brain slices showing a comparison of Cre-dependent RFP expression (red) after the viral vector infusion into the VMH of Nmu-Cre mice, and endogenous ZsGreen1 expression (green) in Nmu-Cre:ZsGreen1 mice. (D) Representative images from coronal brain slices showing neuronal projections from VMH^NMU^ neurons after the viral vector infusion into the VMH of Nmu-Cre mice. Representative images from 1 mouse brain; n = 3 brains were analyzed. Images from C and D were acquired with a confocal microscope with a 10x objective lens. Scale bar: 300 μm. Anatomical abbreviations are listed in Table 5.

**Figure 7.**
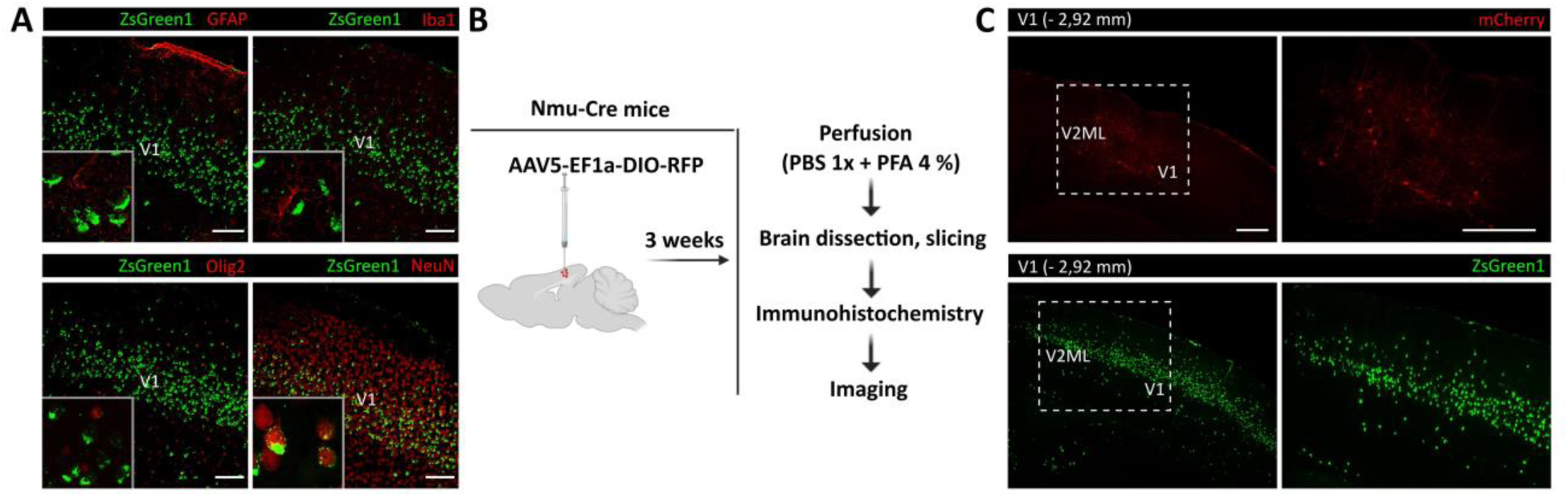
Characterization of NMU neurons in the primary visual cortex (V1) using immunohistochemistry and a viral tracer. (A) Immunohistochemical analysis of brain coronal slices from Nmu-Cre:ZsGreen1 mice at the level of the V1. In green, ZsGreen1 fluorescence; in red, GFAP and Olig2 (secondary antibodies coupled to Cy3), NeuN and Iba1 (secondary antibodies coupled to Cy5). Representative images from 1 mouse brain; n = 3 brains were analyzed. Images were acquired with a confocal microscope at 20x magnification. Areas of higher magnification (40x) are outlined in gray (left-bottom). Scale bar: 100 μm. GFAP: glial fibrillary acidic protein; Iba1: ionized calcium-binding adaptor molecule 1; Olig2: oligodendrocyte transcription factor 2; NeuN: neuronal nuclear protein. (B) Schematic representation of the intracerebral injection of the viral tracer AAV5-EF1a-DIO-RFP into the V1 of Nmu-Cre mice. Three weeks after surgery, mice were transcardially perfused, and brains were processed for immunohistochemical amplification of the RFP fluorescent signal with an anti-mCherry primary antibody and a Cy3-conjugated secondary antibody (red). (C) Representative images from coronal brain slices showing a comparison of Cre-dependent RFP expression (red) after the viral vector infusion into the V1 of Nmu-Cre mice, and endogenous ZsGreen1 expression (green) in Nmu-Cre:ZsGreen1 mice. Areas of higher magnification are indicated by a dashed white box. Representative images from 1 mouse brain; n = 2 brains were analyzed. Images were acquired with a confocal microscope. Scale bar: 300 μm. Anatomical abbreviations are listed in Table 5.

### 3.6. Neuroanatomical characterization of NMU-expressing cells in the VMH and visual cortex

Consistent with literature^26^, we found robust expression of ZsGreen1 in the VMH, consistently expressing moderate to high levels of *Nmu* transcript. Therefore, we selected the VMH to further characterize NMU-expressing neurons (VMH^NMU^). We performed intracerebral injections of the double-floxed AAV5-EF1a-DIO-RFP viral tracer into the VMH of Nmu-Cre mice, to drive Cre-dependent expression of the red fluorescent protein (RFP) (Figure 6B). By infusing the viral tracer into the VMH, RFP expression into the target region allowed us to further confirm that Cre recombinase is indeed expressed and still active in the VMH of adult Nmu-Cre mice (Figure 6C). In addition, taking advantage of the anterograde and retrograde axonal transport ability reported for the AAV5 serotype^60–62^, we described input regions to the VMH^NMU^ (Figure 6C) and output regions receiving projections from the VMH^NMU^ (Figure 6D). A detailed analysis of RFP expression showed RFP-marked cell bodies in the LH and MeA, and in posterior hypothalamic and midbrain regions such as the MM, PH, and PBP, and to a lesser extent in the EW and RLi (Figure 6C), regions where ZsGreen1 expression was described. Terminals from the VMH^NMU^ were found in anterior regions, at the level of the LS and ST, and within the hypothalamus, including the MPA, anterior hypothalamic area (AHA), LH, DMH and paraventricular hypothalamic nucleus (PVH). Projections were also found to the central amygdaloid nucleus (CeA), and to posterior regions including the PAG and, to a lesser extent, the EW (Figure 6D). In addition to the VMH, a clear cluster of ZsGreen1-expressing cells was observed in the visual cortex. This is in contrast with literature^10,21,26^ and our ISH data, that showed a few cells expressing low amount of *Nmu* transcript within the visual cortex. To test whether the *Nmu* promoter was still active in the adult visual cortex, AAV5-EF1a-DIO-RFP was infused in the V1 (Figure 7B), revealing RFP-marked cells in the target region (Figure 7C). The RFP expression pattern was similar to that described for ZsGreen1; however, only a few RFP-marked cells were observed, in contrast with the strong ZsGreen1 signal but in line with the mRNA ISH results obtained in visual areas.

### 3.7. Evaluation of the presence of fast-acting neurotransmitters in NMU neurons

To further characterize VMH^NMU^ neurons, we addressed co-expression of *Nmu* mRNA with markers of the primary excitatory and inhibitory neurotransmitters of the CNS, glutamate and gamma-aminobutyric acid (GABA). Single-molecule fluorescence ISH by RNAscope^®^ was used in coronal brain slices from Nmu-Cre mice for the examination of the targeted RNA within intact cells in the VMH, a region that showed robust *Nmu* mRNA expression. Indeed, fluorescent signal revealing *Nmu* mRNA was found in cells in the VMH, mainly in but also away from the soma, in axons and/or dendrites (Figure 8B) but was not found in the substantia nigra (SN) (Figure 8A), a region where, consistent with previous literature^10,21,26^, neither ZsGreen1 nor *Nmu* mRNA by chromogenic RNAscope^®^ was detected. In the VMH, expressing mainly glutamatergic cells, *Nmu* mRNA signal was found to overlap with *Slc17a6*, the gene encoding for the vesicular glutamate transporter (VGLU2) but not with *Slc32a1* encoding VIAAT (Figure 8B). However, since the observed expression of *Nmu* mRNA was not restricted to the soma of cells in the VMH, *Nmu* expression in GABAergic neurons cannot be excluded. Additionally, we addressed possible co-expression of *Nmu* mRNA and transcripts of the fast-acting neurotransmitters in another region expressing higher number of GABAergic neurons, such as the anterior BNST. In the BNST, that contains mostly GABAergic neurons with a minority of scattered glutamatergic neurons, *Nmu* mRNA was found in both glutamatergic cells and in cells expressing the gene encoding for the vesicular inhibitory amino acid transporter VIAAT (*Slc32a1*) (Suppl. Figure 3). Overall, these results suggest that NMU neurons are not obligate peptidergic neurons, but co-express fast-acting neurotransmitters.

**Figure 8.**
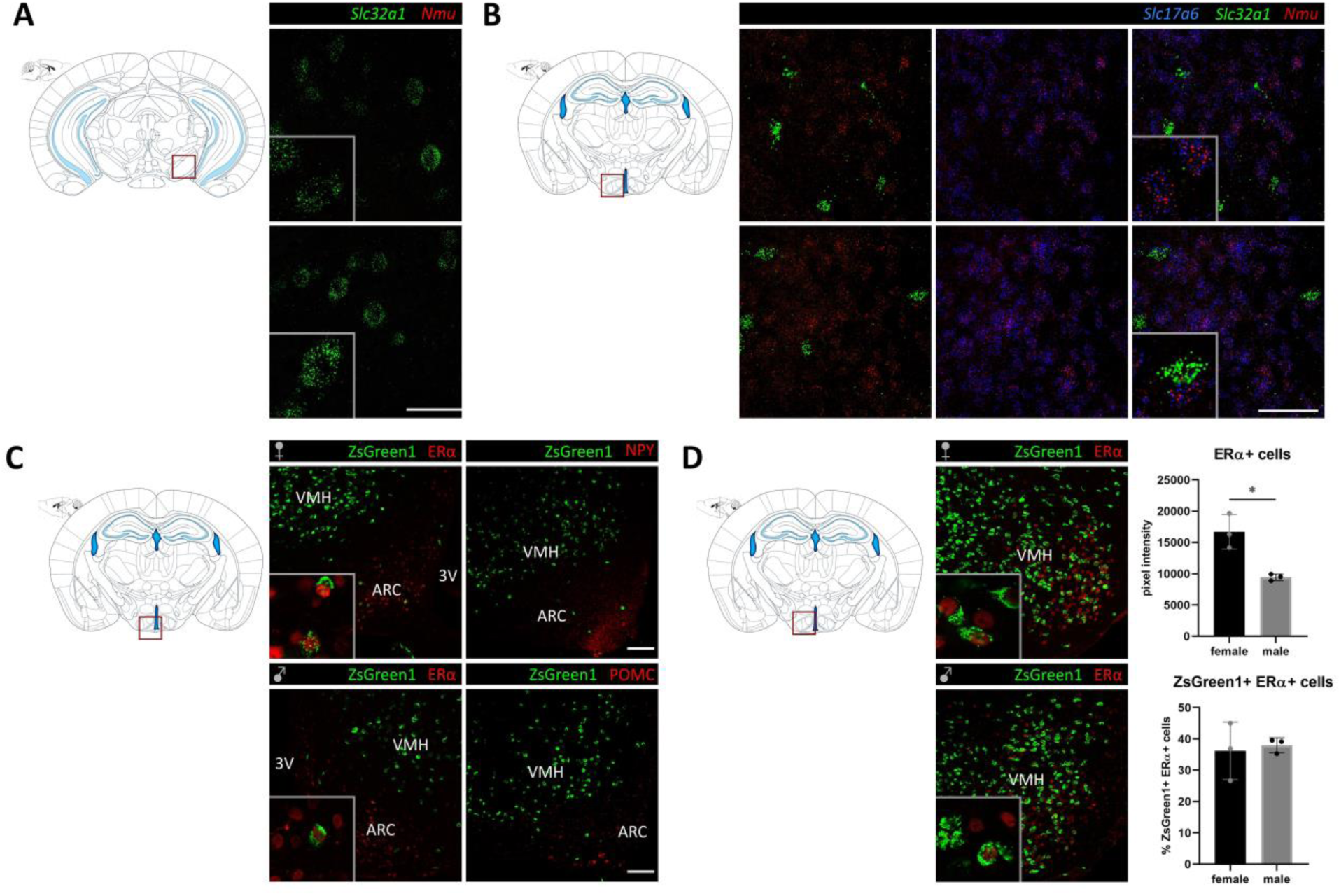
Characterization of NMU neurons in the ventromedial hypothalamic nucleus (VMH) using immunohistochemistry and single molecule fluorescent *in situ* hybridization (RNAscope^®^). (A) RNAscope^®^ for *Slc32a1* (green, VIAAT) and *Nmu* (red) mRNA in the substantia nigra pars compacta (SNc) showing expression of the vesicular transporter VIAAT but failing to detect *Nmu* mRNA. (B) RNAscope^®^ for *Slc17a6* (blue, VGLU2), *Slc32a1* (green, VIAAT) and *Nmu* (red) mRNA in the VMH. Representative images from 1 mouse brain; n = 2 brains were analyzed. Images were acquired with a confocal microscope at 40x magnification. Areas of higher magnification are outlined in gray (left-bottom). Scale bar: 50 μm. (C) Analysis of ZsGreen1 co-expression ERα. Immunohistochemical analysis of brain coronal slices from Nmu-Cre:ZsGreen1 male and female mice at the level of the VMH. In green, endogenous ZsGreen1 fluorescence; in red, ERα (secondary antibody coupled to Cy5). Overlap of ZsGreen1 and ERα in the ventrolateral VMH was quantified using ImageJ (n = 3 males and n = 3 females). Graphs indicating ERα signal measured as pixel intensity (top) and % of ZsGreen1 positive cells expressing ERα (bottom). Data was analyzed by student’s t-test; *p<0,05. (D) Analysis of ZsGreen1 co-expression with ERα, NPY and POMC. Immunohistochemical analysis of brain coronal slices from Nmu-Cre:ZsGreen1 male and female mice at the level of the VMH and arcuate nucleus (ARC). In green, endogenous ZsGreen1 fluorescence; in red, ERα, NPY and POMC (secondary antibody coupled to Cy5). Images of C and D were acquired with a confocal microscope at 20x magnification. Areas of higher magnification (40x) are outlined in gray (left-bottom). Scale bar: 100 μm. Schematic representations of coronal slices with the region of interest outlined in red are depicted for A-D. VIAAT: vesicular inhibitory amino acid transporter; VGLU2: vesicular glutamate transporter 2; Nmu: neuromedin U; ERα: estrogen receptor alpha; NPY: neuropeptide Y; POMC: proopiomelanocortin. Anatomical abbreviations are listed in Table 5.

### 3.8. Evaluation of ZsGreen1 co-expression with specific markers in the VMH and ARC

Additional immunofluorescence analysis was performed to further characterize VMH^NMU^ neurons in coronal brain slices from Nmu-Cre:ZsGreen1 mice (Figure 8C-D). Robust expression of ZsGreen1 was observed in the VMH, while only a few marked cells were shown in the ARC. This expression was consistent with the ISH data in Nmu-Cre mice. Antibodies against pro-opiomelanocortin (POMC) and neuropeptide-Y (NPY) were used as markers of the ARC, while estrogen receptor alpha (ERα) was used as a marker of both the ARC and the VMH. Interestingly, the analysis revealed no overlap with POMC-expressing cells nor NPY-expressing cells (Figure 8C), while less than 40% of ZsGreen1 cells colocalized with ERα in the ventrolateral VMH (vlVMH) of both male and female adult mice (Figure 8D). These results suggest that NMU neurons mainly constitute a distinct population of hypothalamic cells.

## 4. Discussion

In the present study, we generated a Nmu-Cre knock-in mouse model constitutively expressing Cre recombinase in all NMU-producing cells. We validated the model and carried out a complete mapping of NMU expression in both male and female adult mouse brain. Additionally, we further characterized NMU neurons in the VMH using a viral tracer, IHC and ISH analysis.

The strategy that was used to generate the Nmu-Cre knock-in mouse model was based on the absence of regulatory elements that may be impacted by the insertion of the Cre-IRES-*Nmu*-hGHpolyA recombination cassette. Selecting the hypothalamus as the NMU-expressing region, as consistently reported in rodents and humans^10,16,19,21,25,26,32^ and in agreement with this study, our RT-qPCR analysis indicated that the strategy chosen to generate the knock-in mouse model was not interfering with endogenous *Nmu* expression. Our data also highlighted the regional differences reported previously in the literature^1^, comparing the transcript levels between the hypothalamus and the hippocampus. These results were consistent with the clear ZsGreen1 regional differences observed in Nmu-Cre:ZsGreen1 mice, and thus supporting the validation of the model.

Gene targeting is considered the most reliable approach to faithfully mimic endogenous gene expression^38^, with the binary system Cre/LoxP being one of the most extensively used strategies for cell-type-specific genetic targeting^38,40,41^. The popularity of this strategy is reflected in the fact that currently anatomical information is available for more than 200 Cre driver lines in the Transgenic characterization database from the Allen Institute for Brain Science^63^. When using Cre driver lines, and after verifying that the genetic strategy does not interfere with the endogenous expression of the target gene, the main question is how faithfully Cre expression represents endogenous gene expression patterns. First, one of the main strategies to demonstrate that Cre recombination occurs in a Cre mouse line is crossing it with a reporter line that carries a Cre-dependent fluorescent reporter gene. The fidelity of Cre expression can be confirmed by comparing the reporter levels with existing literature and, directly, with the protein and/or mRNA expression of the gene of interest. However, validation of our Nmu-Cre knock-in mouse model was challenging and hampered by the lack of detailed neuroanatomical information in mouse available in the literature, the lack of suitable anti-NMU antibodies and the poor sensitivity of classical mRNA ISH techniques. As such, our present study can be seen as a pioneering study performed with state-of-the-art technologies that brings new insights to the field about the NMU population of neurons in the brain.

In general, our results suggest that the pattern of the observed ZsGreen1 signal broadly represents the endogenous NMU expression. However, the intensity and extent of the transcript signal did not always correlate with the ZsGreen1 signal. ZsGreen1 and *Nmu* transcript expression levels were consistent in several BNST divisions and within the hypothalamic, midbrain, hindbrain and medullary regions analyzed. Our data is in agreement with previous literature specifically in certain hypothalamic (SCh, MPA, VMH, MM) and midbrain (EW) regions^10,21,26^. The SCh is the main region where moderate expression of NMU has been consistently described in both rats and mice^21,26,29^. Our results confirm expression of NMU in this region, in line with the suggested role of the peptide in the regulation of the circadian oscillator system^4^. Remarkably, the neuropeptide neuromedin S (NMS), whose precursor shares high structural similarity with that of NMU, is considered another endogenous ligand of NMUR1 and NMUR2 and has a more restricted expression in the SCh^64^. Indeed, both NMU and NMS have been implicated in the regulation of circadian rhythms^65^. Interestingly, a diurnal rhythm has also been described in the EW^66^, supporting the role of NMU in the regulation of the circadian clock. We also found low to moderate levels of both ZsGreen1 and *Nmu* mRNA in the DMH, a region where previous studies showed discrepancies in terms of transcript expression levels^21,26^. In the ARC, we similarly detected very few ZsGreen1 cells and low *Nmu* mRNA levels, which contrasts with the moderate *Nmu* mRNA expression reported in mice by Graham and colleagues^26^. However, probe sensitivity could be a possible explanation for these discrepancies.

We observed some inconsistencies between ZsGreen1 and *Nmu* transcript levels. We detected very low transcript levels of *Nmu* but high ZsGreen1 signal in the cortex (notably the V1 and V2), thalamus (especially in the reuniens thalamic nucleus [Re] and the anteromedial thalamic nucleus [AM]), PAG and MeA. Funes and colleagues^10^ used RT-qPCR and reported low levels of *Nmu* transcript in the thalamus, similarly to our ISH data. To the best of our knowledge, neither NMU nor *Nmu* mRNA were previously reported in the mouse cortex, PAG or MeA. Interestingly, cortical NMU expression has been described in humans^67^, but expression in rats is still under debate. One study found NMU-LI in pyramidal cells of the cerebral cortex located in the layer V of the somatosensory and motor area^20^. However, these results differed from the study of Ballesta et al.^16^, who did not detect immunoreactive fibers or terminals in the telencephalon. In our present study, the infusion of an AAV vector specifically driving Cre-dependent expression of the fluorescent protein RFP into the visual cortex showed the presence of very few RFP-marked cells. This confirmed our ISH data and indicated that there is indeed Cre expression in the adult cortex, but in very low levels. Interestingly, transgenic overexpression of NMU in mice showed ubiquitous *Nmu* mRNA expression levels compared to wild-type mice, including remarkable expression throughout the cortex^68^. These results support our data showing cortical expression of NMU, pointing to the need to discuss the biological relevance in terms of amount and role of NMU protein in cortical areas under physiological conditions in adult life.

It is not unusual to find the expression pattern of Cre and Cre-reporters more widespread than initially expected^69,70^. In our study, the lack of correlation between the moderate to high ZsGreen1 expression consistently reported throughout the cortex, in certain thalamic and amygdaloid regions, and the very low *Nmu* mRNA levels observed in those regions may be due to a combination of events. First, transient Cre expression during development must be considered. Many transgenes are sensitive to epigenetic regulation and seem to be transiently expressed in cells during the developmental stage but are no longer expressed in adult life. Second, NMU may similarly be expressed in certain regions at early stages of development but silenced at later stages. To our knowledge, no detailed information has been published on the time course of NMU expression in the CNS at different stages of development, but this possibility needs to be considered. Third, even though knock-in mice bypass the positional effect due to random integration, potential ectopic expression of Cre must be contemplated. In our present study, transient expression of NMU and Cre at early developmental stages could result in unanticipated ZsGreen1 expression in offspring of the heterozygous Nmu-Cre and homozygous Ai6 cross, in cells that no longer express NMU in the adult life. ZsGreen1 half-life has been suggested to be at least 26 h in mammalian cells or slightly longer due to its tetrameric structure, in contrast with the monomeric configuration of other well-known fluorescent proteins^58,71^. Thus, a significant accumulation of fluorescent signal until adult life is not expected. In turn, ectopic expression of Cre could result in unanticipated reporter expression. It is known that the efficiency of the recombination depends, in part, on the location of the LoxP sites in the genome and the distance separating them^72,73^. The distance between both LoxP sites in the Ai6 mice may lead to potential spontaneous recombination, resulting in ZsGreen1 expression in cells that do not express Cre recombinase^41^. However, since the unanticipated expression due to spontaneous recombination is expected to be variable in offspring from the same line, several mice need to be examined in order to identify the effects of these unwanted recombination events^70^. In our study, ZsGreen1 signal was analyzed in several male and female adult mice, showing consistent results. It is also important to highlight that every floxed allele can have a different sensitivity to Cre-mediated recombination; this means that different reporter lines can show different Cre activity^72^. Therefore, it may be interesting to cross the new Cre recombinase line with a different reporter line, and to examine in detail reporter expression in certain areas at different developmental stages. Ultimately, even though a significant expression of Cre in adult life due to transient and ectopic expression is not expected, the resulting low Cre expression may be sufficient to induce recombination events between LoxP sites, thus generating unintended, but strong and consistent fluorescent signal^70^. Overall, our results suggest that the ZsGreen1 reporter is indeed a useful tool that reveals which regions express NMU in the brain, but these results cannot be extrapolated to the amount of protein that is actually expressed.

Reporter expression is considered to reflect the end result of the complex modulatory activities of transcription and translation. In addition, the lack of suitable anti-NMU antibodies prevents a direct detection of the NMU protein, hampering a comparison with ZsGreen1 and *Nmu* mRNA. It is important to highlight that in the present study we did not examine *Nmu* mRNA expression across different time points. Certainly, *Nmu* mRNA expression was shown to be differentially modulated in different conditions. Like many other neuropeptides, *Nmu* mRNA expression also exhibits a circadian rhythm in the SCh. In rats, *Nmu* mRNA levels peak between circadian time 4 to 8, which represents the subjective day^29^ and have also been shown to fluctuate during post-natal maturation and oestrus cycle^2^. Therefore, it may be interesting to use a short half-life Cre reporter mice for a longitudinal monitoring of the *Nmu* transcriptional activity in future work. Even though the correlation between mRNA expression and protein expression levels can be relatively low, the detection of the mRNA is generally considered a good indicator of the presence of a protein^74^, and a better approximation of the amount of protein present than the reporter expression. Overall, the NMU distribution revealed by our ZsGreen1 data and ISH results confirms and significantly extends the current knowledge on NMU distribution in the mouse brain.

The description of the distribution of ZsGreen1-expressing cells was performed not only in discrete coronal slices but in the whole brain. The combination of optical tissue clearing and LSFM enabled us to reach a better understanding of the neuroanatomical distribution of NMU-expressing cells in the adult mouse brain, since it provides a complete picture in intact tissue. The protocol optimized in this study represents a novel option for tissue clearing, combining the powerful and effective CUBIC clearing protocol with the complementary refractive index matching of CUBIC-R and ECi to obtain optimally optical cleared tissue. The analysis unveiled a continuous population of ZsGreen1-expressing cells extending from the ventral forebrain to the caudal midbrain, and beyond the midbrain through the pons and the medulla. It is tempting to speculate on the existence of a midline circuit of NMU-expressing regions projecting to and receiving inputs from other NMU-expressing regions. Indeed, the results of the infusion of the double-floxed AAV5-EF1a-DIO-RFP viral tracer into the VMH suggest this possibility. The bidirectional axonal transport ability of the AAV5 serotype has been observed when injected into the CNS in both mouse and rat^60–62^, while it did not exhibit anterograde transneuronal transport^75^. Thus, the detection of marked cell bodies in distant areas from the target region strongly suggest retrograde transport of the viral tracer and transduction of NMU-expressing neurons. Our results revealed the existence of NMU-expressing neurons in hypothalamic, amygdaloid and midbrain regions projecting to the VMH, and VMH^NMU^ neurons projecting to anterior and posterior areas, from the LS and ST to the PAG and EW. The connectivity reported here partially overlaps with the connectional architecture reported for vlVMH^Esr1^ neurons^76^, including projections to the BNST and hypothalamic (AHA, MPA, DMH), thalamic (PVH) and midbrain (PAG) regions. Interestingly, we found a substantial amount of NMU neurons in the vlVMH overlapping with ERα, supporting the hypothesis that NMU plays a modulatory role in reproduction and energy expenditure^2,77,78^. However, NMU neurons were found not only in the vlVMH, but throughout all three subdivisions of the VMH, in contrast with many other neuronal subpopulations identified in this nucleus, such as the aforementioned ERα, leptin receptor, pituitary adenylate cyclase-activating peptide (PACAP) or cholecystokinin receptor B (CCKBR), among others^79,80^. The expression pattern of NMU neurons in the VMH and ARC, the limited colocalization with ERα in both regions, and the absence of colocalization with other well-described hypothalamic markers, such as POMC and NPY, suggest that NMU neurons constitute mainly a unique population of hypothalamic cells. Overall, the expression of NMU in the VMH and the connectional pattern to and from VMH^NMU^ supports the idea of a midline NMU circuit with the VMH as a key node. In that sense, there is a relevant overlap between neuroendocrine functions attributed to the VMH and those described for NMU. The VMH is considered an integral region in many neuroendocrine functions, such as metabolic regulation and energy homeostasis, thermoregulation, bone remodeling as well as stress and anxiety related behaviors^81,82^, functions that have also been described for NMU^4^. A central stress-integrative system involving forebrain, amygdaloid, hypothalamic and midbrain region has been reported^83^, pointing out a role of the VMH and PH outputs to the PVH in the regulation of the stress response. The suggested NMU midbrain circuit comprise most of those regions; certainly, a role of NMU in the modulation of the stress response in rodents has been extensively addressed^29,84–90^. It may be interesting to perform a detailed analysis of the connectivity between these NMU-expressing regions to gain insights into the connectional architecture of this potential NMU circuit. In addition, the presence of RFP marked cells in hypothalamic and midbrain regions constitute an additional validation of the mouse model, since these results confirmed Cre activity in the adult life in regions where the reporter ZsGreen1 protein and *Nmu* mRNA were detected. Furthermore, the specific targeting of NMU neurons with the double-floxed viral tracer opens the door to the use of chemogenetic and optogenetic interventions in Nmu-Cre mice to modulate specific sub-populations of NMU-expressing neurons. This may allow further exploration of the role of NMU in the mouse brain.

In contrast to classic neurotransmitters, neuropeptides were initially considered as transmitters of a modulatory nature^91^. Nowadays, it is well known that neuropeptides are signaling molecules widely distributed throughout the CNS where they commonly occur with, and are complementary to, classic neurotransmitters, controlling a wide range of behaviors^92^. However, the idea of obligate peptidergic neurons has also recently been addressed in the mouse brain^93^. Our results showing co-expression of transcripts for fast-acting neurotransmitters and *Nmu* mRNA in VMH^NMU^ neurons are consistent with the co-transmission described for many hypothalamic peptidergic neurons^92,94^. This co-expression was also found in another NMU-expressing region, the BNST. The effects of NMU are linked to two GPCRs, driving delayed changes in neuronal activity through a broad spectrum of signaling pathways. Although the functional effects of the co-transmission of neuropeptides and fast-acting neurotransmitters are still not fully understood^95^, this phenomenon potentially gives NMU a much broader and diverse range of biological effects^92,96,97^. In addition, our data showed expression of the *Nmu* transcript following a punctate pattern not confined to perinuclear somatic regions, suggesting that local protein synthesis may play a role^98,99^.

Taken together, our results strongly suggest that Cre expression in the Nmu-Cre knock-in mouse model largely and faithfully reflects NMU expression in the adult mouse brain, without altering endogenous *Nmu* expression. In addition, we provided a detailed whole-brain mapping of NMU-expressing neurons. Our results revealed a potential midline NMU modulatory circuit with the VMH as a key node. We believe that this mouse model can serve as a useful tool for future investigations on NMU neurons in the CNS and the periphery.

## Abbreviations

AAV: Adeno-associated virus
*B2m*: beta-2 microglobulin
BAC: Bacterial artificial chromosome
cDNA: Complementary DNA
CNS: Central nervous system
CUBIC: Clear, unobstructed brain imaging cocktails and computational analysis
DAPI: 4’,6-diamidino-2-phenylindole dihydrochloride
ECi: Ethyl cinnamate
EGFP: Enhanced green fluorescent protein
ERα: Estrogen receptor alpha
ES: Embryonic stem cells
FRT: Flippase recognition target
GABA: Gamma-aminobutyric acid
GFAP: Glial fibrillary acidic protein
*Hprt1*: Hypoxanthine guanine phosphorybosyl transferase 1
Iba1: Ionized calcium-binding adaptor molecule 1
IHC: Immunohistochemistry
IRES: Internal ribosome entry site
ISH: *In situ* hybridization
LSFM: Light-sheet fluorescence microscopy
NBF: Neutral-buffered formalin
NeuN: Neuronal nuclear protein
NMS: Neuromedin S
NMU: Neuromedin U
NMU-LI: NMU-like immunoreactivity
NPY: Neuropeptide Y
Olig2: Oligodendrocyte transcription factor 2
PBS: Phosphate-buffered saline
PFA: Paraformaldehyde
POMC: Proopiomelanocortin
RFP: Red fluorescent protein
RIA: Radioimmunoassay
RT-qPCR: Quantitative reverse-transcription polymerase chain reaction
TB: Tris buffer
TBS: Tris-buffered saline
TBS-T: Tris-buffered saline-Triton-X
VIAAT: Vesicular inhibitory amino acid transporter
VGLU2: Vesicular glutamate transporter 2
VMH^NMU^: NMU-expressing neurons in the ventromedial hypothalamic nucleus

## Acknowledgements

We would like to thank Gunter Leuckx for the theoretical support and Anke De Smet, Eddy Himpe, Geoffrey Duqué, Yves Heremans and Astrid Deryckere for the practical assistance.

