## Supplementary material for "Neuroanatomical characterization of the Nmu-Cre knock-in mice reveals an interconnected network of unique neuropeptidergic cells"

**Supplementary figure 1**

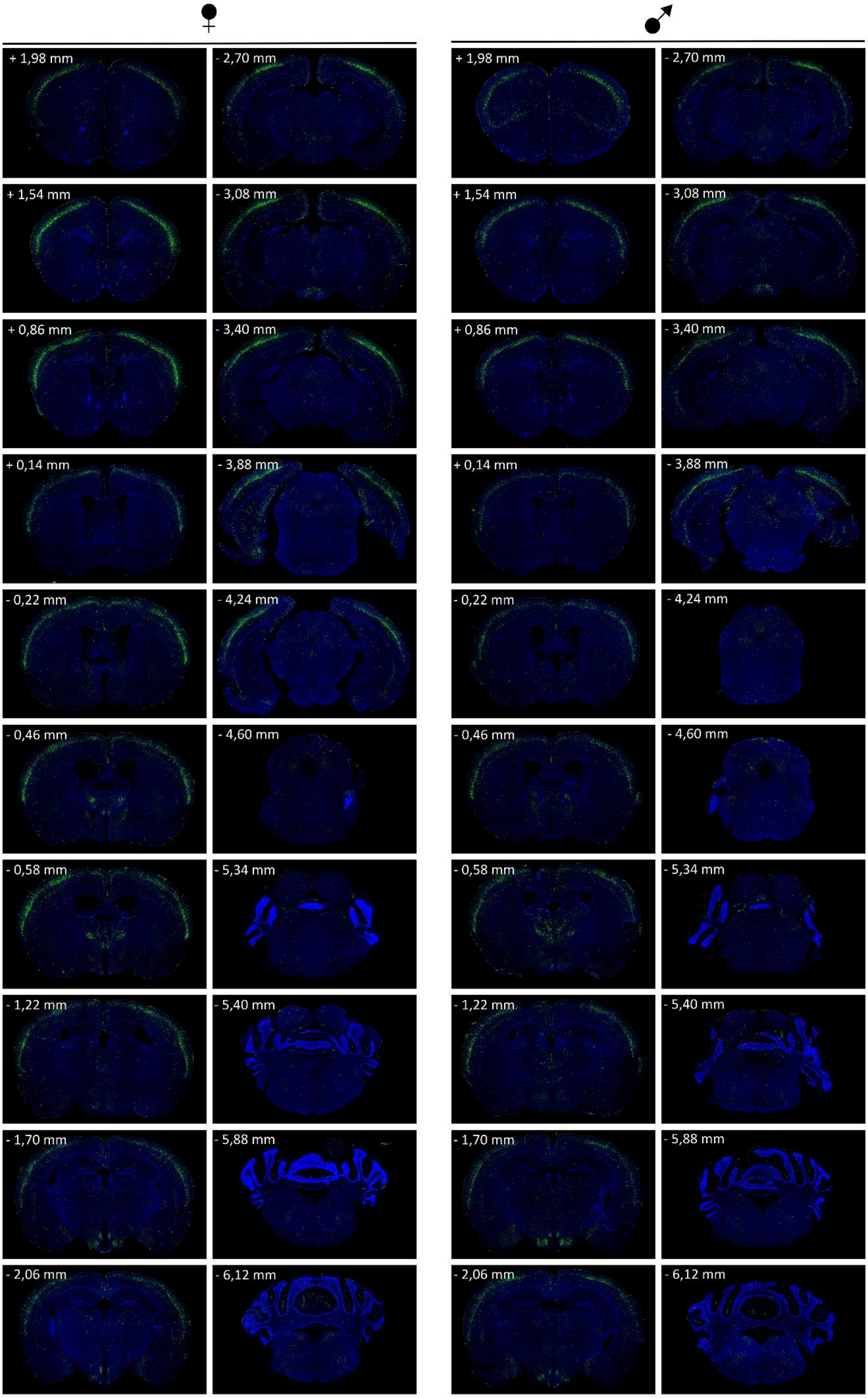

**Supplementary figure 1. Overview of ZsGreen1 expression in coronal brain slices obtained from Nmu-Cre:ZsGreen1 mice**. Representative large sections from a female (left) and a male (right) mouse. DAPI nuclear staining (blue), ZsGreen1 (green). Images were acquired with a fluorescent microscope with a 4x objective lens. Numbers denote neuroanatomical coordinates relative to Bregma.

### Supplementary figure 2

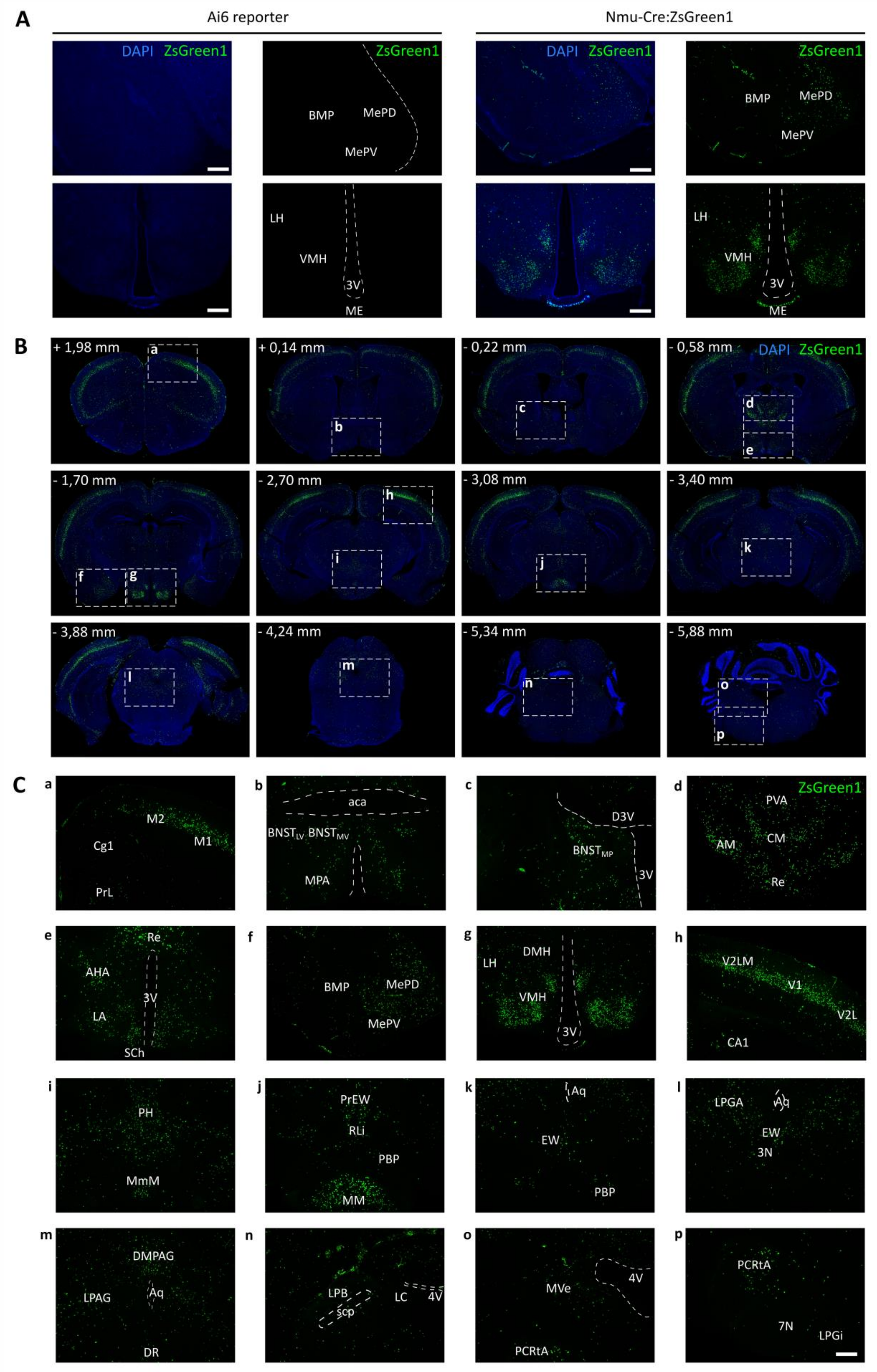

**Supplementary figure 2. Neuroanatomical distribution of ZsGreen1 expression in coronal brain slices from male Nmu-Cre:ZsGreen1 mice.** (A) ZsGreen1 fluorescence background levels detected in coronal slices from Ai6 reporter mice at the level of the amygdala (top) and the ventromedial hypothalamic nucleus (bottom). From left to right, Ai6 merged image with DAPI nuclear staining (blue) and ZsGreen1 (green); Ai6 ZsGreen1 channel; Nmu-Cre:ZsGreen1 merged image with DAPI and ZsGreen1; Nmu-Cre:ZsGreen1 green channel. (B-C) Distribution of ZsGreen1-expressing cells in coronal slices (representative images from 1 male; n = 4 males were analyzed). The qualitative analysis was performed on 25-35 coronal slices per mouse brain by two independent observers. DAPI nuclear staining (blue), ZsGreen1 (green). Areas of higher magnification are indicated by a dashed white box. Images were acquired with a fluorescent microscope with 4x (B) and 10x (C) objective lens. Numbers denote neuroanatomical coordinates relative to Bregma. Scale bar: 300 µm. Anatomical abbreviations are listed in Table 5.

**Supplementary figure 3**

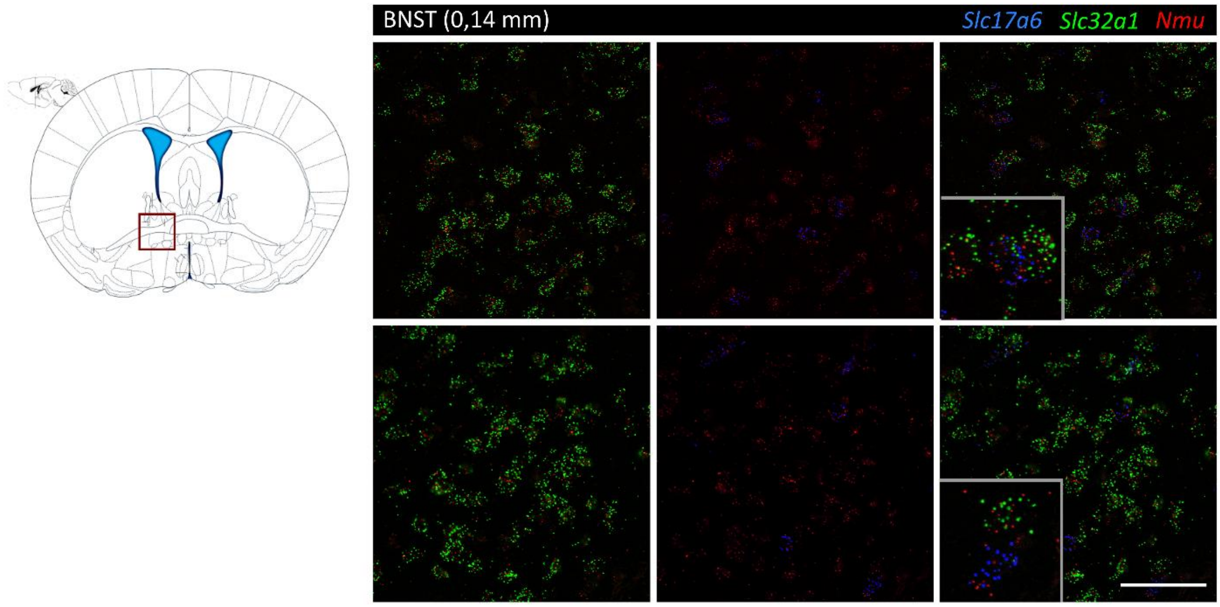


**Supplementary figure 3. Evaluation of the presence of fast-acting neurotransmitters in NMU neurons in the bed nucleus stria terminalis (BNST) using single molecule fluorescent *in situ* hybridization (RNAscope®).** RNAscope® for *Slc17a6* (blue, VGLU2), *Slc32a1* (green, VIAAT) and *Nmu* (red) mRNA in the ST. Representative images from 1 mouse brain; n = 2 brains were analyzed. Images were acquired with a confocal microscope. Areas of higher magnification are outlined in gray (left-bottom). Scale bar: 50 µm. Numbers denote neuroanatomical coordinates relative to Bregma. A schematic representation of a coronal slice with the region of interest outlined in red is depicted. VIAAT: vesicular inhibitory amino acid transporter; VGLU2: vesicular glutamate transporter 2; Nmu: neuromedin U.

**Video 1. Neuroanatomical distribution of ZsGreen1 expression in a cleared mouse brain.** 3D volume rendering of half brain from a representative Nmu-Cre:ZsGreen1 mouse and. Objective 2x, digital zoom 0,63. 488 nm and 561 nm lasers were used to reveal ZsGreen1 endogenous fluorescence (green) and background signal (gray), respectively.
